# Mapping social reward and punishment processing in the human brain: A voxel-based meta-analysis of neuroimaging findings using the Social Incentive Delay task

**DOI:** 10.1101/2020.05.28.121475

**Authors:** D. Martins, L. Rademacher, A. S. Gabay, R. Taylor, J. A. Richey, D. Smith, K. S. Goerlich, L. Nawijn, H.R. Cremers, R. Wilson, S. Bhattacharyya, Y. Paloyelis

**Affiliations:** Department of Neuroimaging, Institute of Psychiatry, Psychology and Neuroscience, King’s College London, De Crespigny Park, London SE5 8AF, UK; Department of Psychiatry and Psychotherapy, University of Lübeck, Lübeck, Germany and Department of Psychology, Goethe University Frankfurt, Germany; Department of Experimental Psychology, University of Oxford, New Radcliffe House, Oxford OX2 6NW, UK; Department of Psychology, Virginia Tech, Blacksburg USA; Department of Psychology, Temple University, Philadelphia, PA, 19122, USA; Department of Biomedical Sciences of Cells & Systems, Section Cognitive Neuroscience, University Medical Center Groningen, University of Groningen, Groningen, The Netherlands; Department of Psychiatry, Amsterdam UMC, Vrije Universiteit Amsterdam, Amsterdam Neuroscience, Amsterdam, the Netherlands; Department of Clinical Psychology, University of Amsterdam, Amsterdam, the Netherlands; Department of Psychosis Studies, Institute of Psychiatry, Psychology and Neuroscience, King’s College London, De Crespigny Park, London

**Keywords:** Social incentive delay, Social Reward, Social Punishment, Anticipation, Feedback, Anisotropic Effect Size Signed Differential Mapping

## Abstract

Social incentives (rewards or punishments) motivate human learning and behaviour, and alterations in the brain circuits involved in the processing social incentives have been linked with several neuropsychiatric disorders. However, questions still remain about the exact neural substrates implicated in social incentive processing. Here, we conducted four Anisotropic Effect Size Signed Differential Mapping voxel-based meta-analyses of fMRI studies investigating the neural correlates of the anticipation and receipt of social rewards and punishments using the Social Incentive Delay task. We map the regions involved in each of these four processes in the human brain, identify decreases in the BOLD signal during the anticipation of both social reward and punishment avoidance that were missed in individual studies due to a lack of power, and characterise the effect size and direction of changes in the BOLD signal for each brain area. Our results provide a better understanding of the brain circuitry involved in social incentive processing and can inform hypotheses about potentially disrupted brain areas linked with dysfunctional social incentive processing during disease.

**Highlights:**

- Voxel-based meta-analysis of the neural underpinnings of social incentive processing
- We map the brain regions involved in the processing of social incentives in humans
- We identify new regions missed in individual studies as a result of lack of power
- Our work can inform research on pathological brain processing of social incentives

## Introduction

Social incentives (rewards or punishments) are crucial for learning and adaptive behaviour (Fehr and Camerer, 2007). Disruption of the brain circuits processing social incentives has been suggested to be at the heart of several neuropsychiatric disorders characterised by dysfunction in social interactions, such as autism spectrum disorder (Kohls et al., 2013), psychosis (Radua et al., 2015) or mood disorders (Naranjo et al., 2001). However, questions still remain about the exact neural circuitries underlying the processing of social incentives in the human brain.

The neural mechanisms prompting an organism to approach/avoid a potential reward/punishment have been extensively explored in several studies using monetary incentives (for an overview see (Dugre et al., 2018a; Oldham et al., 2018a; Wilson et al., 2018)). Among the various paradigms available, the monetary incentive delay (MID) task is among the most widely-used ones. Performance-dependent reward and punishment processing can be divided in two distinct temporal phases: an anticipation phase, where the prospect of a reward/loss is initially encountered, and an outcome phase (also called the receipt or consummatory phase), where the reward/loss is received or omitted (Oldham et al., 2018a). In the classical MID task, participants are asked to perform a simple motor reaction time task (i.e. press a button while a target is on the screen) in various incentive contexts defined by the presentation of cues indicating one of a range of positive or negative monetary rewards or punishment that may follow the response to the target. The cue is followed by a short delay period (the anticipation phase), before the target appears. Participants are instructed to respond to the target as quickly as possible, and the success rate is set to a fixed value (typically at 67% of the trials) by adjusting the duration of the target presentation trial by trial and for each individual separately. Finally, in the outcome/feedback phase, the performance-dependent outcome is presented (attainment or loss of a potential monetary gain, or no monetary outcome in neutral trials). The combination of the MID task design with functional magnetic resonance imaging (fMRI) offers the opportunity to assess the neural signatures specifically associated with the anticipation or receipt/avoidance of monetary wins/losses. Recent meta-analyses of fMRI studies using the MID task have identified several areas consistently involved in the anticipation and/or receipt of monetary rewards/punishments (Dugre et al., 2018a; Oldham et al., 2018a; Wilson et al., 2018). These meta-analyses have found considerable overlaps in the networks activated during monetary reward and loss anticipation, including the dorsal and ventral striatum, amygdala, insula and supplementary motor cortex. They have also identified neural networks engaged by monetary reward receipt — including the ventral striatum, the orbitofrontal cortex/ventromedial prefrontal cortex, amygdala, and the posterior cingulate.

Building on findings from the MID task, some studies have inspected if the same neural circuits that underpin the processing of monetary incentives also underpin the processing of social incentives. Studies have investigated the neural underpinnings of the anticipation and receipt of social rewards/punishments using a variant of the MID task known as the Social or Affective Incentive Delay task – henceforth referred to as SID task (for an overview see (Gu et al., 2019). The SID task shares a similar structure as the MID task, but uses social stimuli that are inherently reinforcing or punishing (i.e. smiling or angry faces, positive/negative verbal messages, thumbs-up or down gestures) as incentives. Fewer studies have used the SID task than those using the MID task though. Individually, studies using the SID task have started to delineate the brain regions involved in the processing of social rewards/punishments (Gu et al., 2019). On the one hand, there is evidence that the processing of social incentives frequently evokes the activity of generic brain regions involved in reward prediction and value encoding, such as the striatum and the orbitofrontal cortex (Barman et al., 2015b; Rademacher et al., 2014; Spreckelmeyer et al., 2009). On the other hand, some studies have also suggested that the processing of social incentives might involve additional brain regions thought to be critical for social-cognitive processes, such as the temporoparietal junction, dorsomedial prefrontal cortex, precuneus, and superior temporal gyrus(Barman et al., 2015b; Goerlich et al., 2017b; Spreckelmeyer et al., 2013b). However, a clear picture of the brain circuits underpinning social reward and punishment processing in the human brain is still missing due to inconsistencies in the areas identified across individual studies. This lack of consistency makes it difficult to inform targeted hypotheses about potentially disrupted areas in neuropsychiatric disorders, based on our understanding of the neural underpinnings of social reward/punishment processing in healthy individuals (Cremers et al., 2015; Delmonte et al., 2012; Kohls et al., 2018).

Meta-analyses can provide valuable help in our efforts to delineate a precise and fine-grained characterization of the neural substrates involved in both the anticipation and receipt phases of social incentive processing. They can help address one major limitation of individual studies, namely the lack of sufficient power due to small sample sizes. Indeed, lack of statistical power is a well-documented phenomenon in neuroimaging studies (Cremers et al., 2017; Poldrack et al., 2017) and comes at the cost of increased risks for both type I and type II errors, undermining replicability across independent studies (Button et al., 2013; Cremers et al., 2017; Kim, 2015). Therefore, meta-analyses of neuroimaging studies have become increasingly important (Muller et al., 2018).

To date, there has been only one attempt to quantitatively summarize neuroimaging findings from the SID task (Gu et al., 2019). This meta-analysis focused specifically on the anticipation social rewards (it did not examine the receipt phase of social rewards, or the processing of social punishment at all). Given the role that differences in the processing of social punishments may play in a wide range of neuropsychiatric disorders (such as major depressive disorder (Kumar et al., 2017) or antisocial personality disorder (Gong et al., 2019)), characterizing the neural underpinnings of social punishment processing in healthy samples is paramount to identify how disruption of these circuits might give rise to impairments in social cognition during pathology. Furthermore, this previous study used a method for the meta-analysis of neuroimaging data, Activation Likelihood Estimation (ALE) that, although popular, comes with some important limitations (Radua and Mataix-Cols, 2012). For instance, the ALE method is based on the reported peak coordinates in individual studies. However, reported peak coordinates may be biased as they depend on the statistical significance thresholds that were used, which may be arbitrary, and the power in individual studies, which tends to be less than the commonly accepted standard of at least 80%, as highlighted above (Cremers et al., 2017). Furthermore, an ALE meta-analysis does not produce a statistical measure of effect-size or its variance, or show the direction of the effect, thus making the interpretation of the biological significance of the results challenging. Instead, an ALE meta-analysis provides an informative summary of statistically significant fMRI results across a number of studies in a field, based on the spatial convergence of neuroimaging findings across experiments (Radua and Mataix-Cols, 2012).

Here, we aimed to quantitatively synthetize the evidence from fMRI studies using the SID tasks. We generate a fine-grained characterization of the neural circuitry implicated in the anticipation and receipt of both social rewards and punishments. The method used provides effect sizes and direction of change in the BOLD signal for each brain area identified in the analysis. To achieve this, we used Anisotropic Effect Size Signed Differential Mapping (AES-SDM) (Radua et al., 2012; Radua et al., 2014). AES-SDM can combine reported peak information (coordinates and t-values) from some studies, with original statistical parametric maps (SPMs) from other studies, to produce an estimate of the magnitude and direction of the effect sizes of changes in the BOLD signal. The inclusion of unthresholded original statistical maps helps to address issues of low statistical power by preserving information from voxels that did not reach significance in the original studies because of power limitations. In addition, since AES-SDM is a method based on effect sizes, it preserves information on the sign of the effect (e.g. increases or decreases in the BOLD signal), therefore allowing a more straightforward and biologically plausible interpretation of the results (Radua and Mataix-Cols, 2012).

## 1. Materials and Methods

### 1.1. Literature search

We conducted searches in PubMed, EMBASE and OVID using the terms (“social incentive delay” OR “affective incentive delay” OR “SID” OR “AID”) AND (“fMRI” OR “functional magnetic resonance imaging”) on 3^rd^ May 2018. Our search query was adapted in accordance with the specification of each database. Our search strategy was tailored to include all human task-based fMRI studies conducted with healthy individuals, published up to this date (irrespectively of publication date or age, sex and ethnicity of the subjects), and which were original reports on the neural substrates associated with social reward and punishment anticipation and receipt (as assessed by task-based fMRI using the social/affective incentive delay task).

### 1.2. Study selection

We performed study selection with the help of the reference manager software *Rayyan* (Ouzzani et al., 2016). We imported all results from our literature search into *Rayyan* and started by removing all duplicated studies using the *Rayyan* smart group function for duplicates. Then, two independent reviewers (DM and RT) screened all titles, abstracts and keywords to select eligible references for further scrutiny. Any disagreements were adjudicated by consensus. The two independent reviewers then assessed all potentially eligible full-texts against the inclusion/exclusion criteria to decide on the inclusion/exclusion of the reference. Any disagreements were adjudicated by consensus. The inclusion criteria were: original article; written in English; accepted for publication; full-text available; be a task-based fMRI study using the social/affective incentive delay task; to have included healthy individuals. In the case of studies including clinical samples, only data of healthy controls were considered. In the case of pharmacological studies, only data coming from the placebo arm were included. If it was evident that studies used overlapping samples or the same sample then only one of those studies was used (either the one for which coordinates/statistical maps could be obtained or the one using the biggest sample). Finally, to minimize as much as possible searching bias we screened all reference lists from eligible publications to identify further potentially relevant studies. During the whole process of selection, we recorded reasons for exclusion. A flow chart (i.e. PRISMA type) (Shamseer et al., 2015) was used to trace the overall process (Fig.1).

**Fig. 1.**
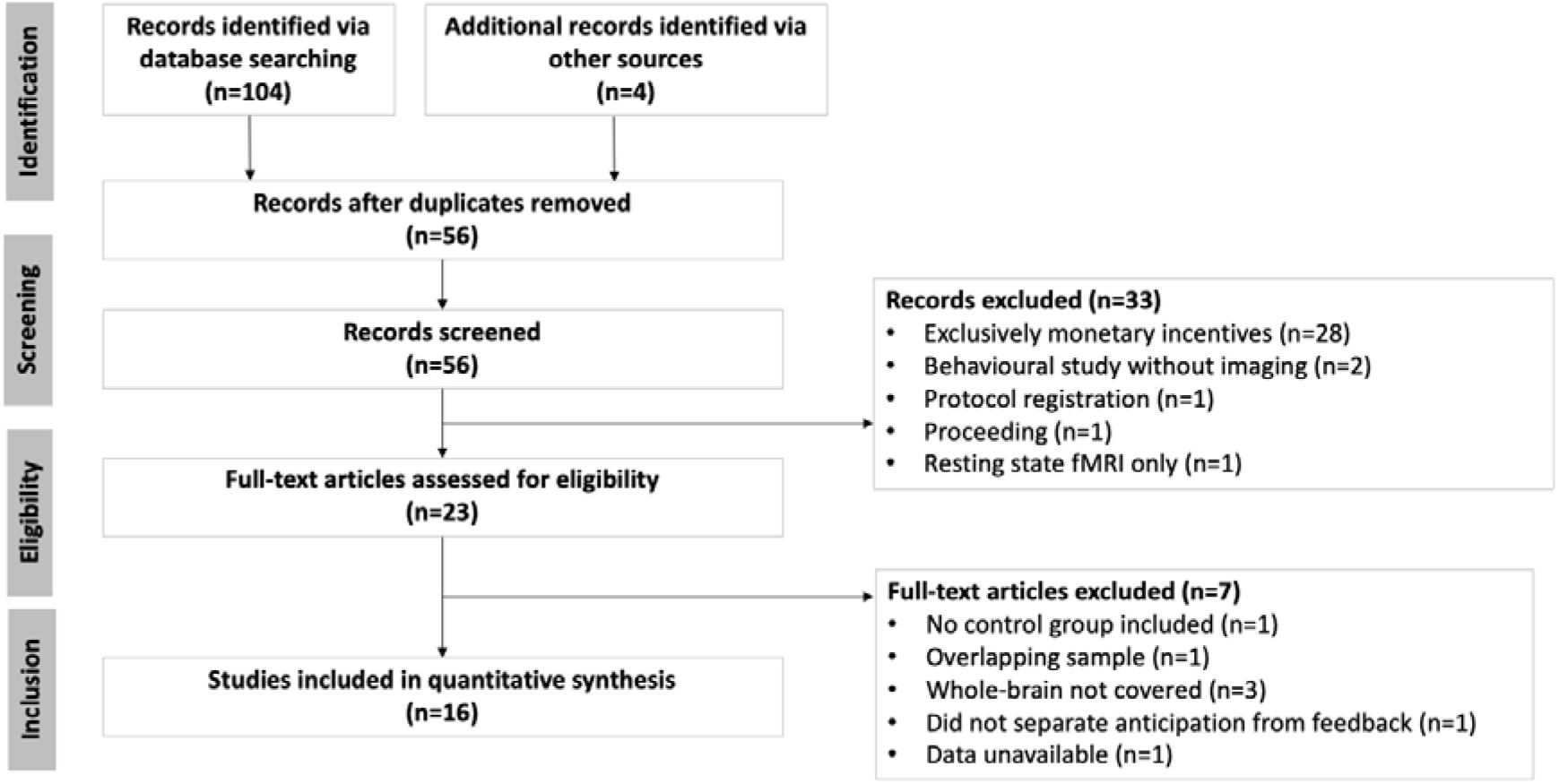
Systematic review process. PRISMA flowchart showing study selection for meta-analysis.

### 1.3. Data extraction

From each included study, two independent reviewers (DM and RT) extracted data on authors, title, year of publication, participants’ gender, age, physical and mental health status, experimental paradigm (including the type and source of social reward/punisher used, the type of neutral comparator, success rate), imaging setup and data analysis method used (whole-brain or region-of-interest, software), and relevant behavioural data (such as hit rates). In respect to the neuroimaging data, we were interested in the following contrasts: anticipation social reward versus anticipation neutral feedback; anticipation social punishment avoidance versus anticipation neutral feedback; receipt social reward versus receipt neutral feedback; receipt social punishment versus receipt neutral feedback. Since one of the main strengths of the meta-analytic method we used here is to allow for the inclusion of statistical maps (which increase the precision and accuracy of the results), we contacted all authors to obtain the statistical maps for these four contrasts. When authors could not provide statistical maps for the contrasts of interest, we used the published peak coordinates and corresponding statistics, if available. When neither was available or could not be obtained from the authors, we excluded the study from further analyses (this resulted in the exclusion of 1 study (Li et al., 2016)). To prevent biasing our meta-analytic maps towards regions identified through the use of more liberal analytic approaches, for the peak coordinates data, we only included studies reporting results from whole-brain analysis (irrespective of the threshold used, provided that the threshold used for each contrast was the same across the whole brain) and excluded studies with partial brain coverage (i.e. only the basal ganglia) or results based on region-of-interest analyses. We also excluded one study which by design did not allow the separation of anticipation from feedback (Kohls et al., 2018).

### 1.4. Meta-analyses

#### Anisotropic Effect Size Signed Differential Mapping

We conducted the meta-analyses using a voxel-wise random effects model implemented in the Anisotropic Effect Size Signed Differential Mapping (AES-SDM) software (www.sdmproject.com). Briefly, AES-SDM is a weighted, voxel based meta-analytic method which has been validated against mega-analyses and used in several functional and structural MRI meta-analyses. It recreates voxel-level maps of Hedge’s *g* effect sizes, their variances and the direction of the effects, and allows the inclusion of both peak information (coordinates and t-values) and statistical parametric maps (Radua et al., 2012). The conversion from t-statistics to Hedge’s *g* is carried out using standard statistical formulae. Hedge’s *g* is equivalent to Cohen’s *d*, corrected for small sample sizes (Radua et al., 2012). When statistics are only available for reported peak coordinates, the effect size is exactly calculated at these peaks and then estimated in the remaining voxels depending on their distance from these peaks, using an un-normalised Gaussian kernel, which is multiplied by the effect size of the peak. The use of effect sizes in the calculation has been shown to increase the accuracy of estimation of the true signal compared to alternative methods (Radua et al., 2012). Additionally, the inclusion of statistical parametric maps has been shown to substantially increase the sensitivity of voxel-based meta-analyses. For example, in the initial validation study of the method, sensitivity increased from 55% to 73% and 87% with the inclusion of just one and two statistical maps, respectively (Radua et al., 2012). The detailed AES-SDM approach has been described elsewhere (Radua et al., 2012; Radua et al., 2014).

We conducted four separate meta-analyses, one for each of our four contrasts of interest. For each of the meta-analyses, meta-analytic effect-sizes were voxel-wise divided by their standard error to obtain AES-SDM z-values. As AES-SDM z-values for each meta-analytic brain map may not follow a standard normal distribution, a null distribution was empirically estimated for each meta-analysis. Specifically, null distributions were obtained from 50 whole brain permutations (which, multiplied by the number of voxels, resulted in about four million values per null distribution); previous simulation work has found that permutation-derived AES-SDM thresholds are already stable with even 5 whole-brain permutations. Voxels with AES-SDM z-values corresponding to p-values <0.001 were considered significant, but voxels with AES-SDM z-values <1, or in clusters with less than 10 voxels, were discarded in order to reduce the false positive rate. While these thresholds do not strictly apply family-wise correction for multiple comparisons, previous empirical validation work of this method has found these parameters to provide optimal sensitivity while maintaining the false positive rate below 5% (Radua et al., 2012).

#### 2.4.2. Sensitivity and sub-group analyses

We assessed the robustness of our findings by conducting Jackknife sensitivity analysis where we iteratively repeated the meta-analysis for each of the four contrasts leaving out one study at a time. In order to synthesize mean Jackknife maps for each contrast and meaningfully interpret them, we first thresholded each individual Jackknife map, for each contrast, using the significance thresholds outlined above, and then binarized them and combined their information into a single overlapping density map of significant voxel-wise data for each contrast. This allowed the visual identification of voxels in terms of density (the higher the density, the more the individual Jackknife maps that a given voxel reached significance), which provides an estimate of replicability across studies. We considered a certain voxel to be robust if it reached significance in more than 75% of the Jackknife leave-one-out meta-analyses, showing that our main findings are not driven by the inclusion of specific studies.

The studies we considered used a variety of methodological approaches. Some studies used verbal feedback while other studies used emotional faces for feedback, some studies used static and others dynamic emotional faces, some studies examined the effects of a pharmacological intervention using a placebo-controlled design (we only considered data form the placebo condition). Therefore, we ran further sensitivity analyses where the number of included studies permitted it (that is, where >5 studies would be included in the respective sub-group meta-analysis). Specifically, for our anticipation and receipt of social reward meta-analyses, we ran meta-analyses on subgroups of studies: 1) including only studies using emotional faces as feedback; 2) including only studies using static emotional faces; 3) including only non-pharmacological studies. By default, we did not conduct any of these sub-group meta-analyses for the anticipation and receipt of social punishment conditions as the number of included studies would be less than 5. However, we did conduct one subgroup meta-analysis for the social punishment avoidance anticipation condition because one study used an implementation of the SID task that was conceptually different from the rest (Nawijn et al., 2017). Specifically, in Nawijn’s et al. implementation of the task, the success rate in the social punishment avoidance condition was set to approximately 34%, making social punishment avoidance the least frequently anticipated outcome (in all other studies, the success rate in the social punishment avoidance condition was set to approximately 66%). The subgroup analysis would allow us to test whether our results were unduly influenced by the inlcusion of Nawijn’s et al. study.

#### 2.4.3. Heterogeneity and publication bias

Heterogeneity is a problem that can arise when undertaking meta-analysis (Muller et al., 2018; Radua and Mataix-Cols, 2012). Ideally, a meta-analysis should combine results from studies using the same procedures and experimental protocols. Heterogeneity refers to the proportion of variability across studies that is due to methodological variability relative to that from sampling error. Heterogeneity can be quantified and provide an index of whether the assumption that all studies use the same procedures and experimental protocols is met (Radua and Mataix-Cols, 2012). When considerable heterogeneity is found, it raises questions about how valid it is to combine all the data in one single meta-analysis. When possible, instances of substantial heterogeneity should be investigated further using subgroup and metaregression analysis (described below) to identify potential factors underlying this heterogeneity. We used the I^2^ statistic maps provided by the software to assess heterogeneity. To identify areas of increased heterogeneity, we thresholded these maps for I^2^>40% (since values of I^2^ < 40 are typically assumed to not constitute important heterogeneity (Higgins et al., 2003)) and masked them to retain only voxels where we found significant increases/decreases in BOLD in the main meta-analyses. Voxels with I^2^ between 40 and 60 indicate areas of moderate heterogeneity, between 60 and 90 substantial heterogeneity and higher than 90 strong heterogeneity(Higgins et al., 2003).

In turn, publication bias occurs when the outcome of an experiment or research study influences the decision to publish it (Joober et al., 2012). Increased publication bias means that a study is less likely to be published if the findings are null. We used funnel plots and the Egger test as implemented by the software to examine publication bias (Lin and Chu, 2018). Briefly, effect size estimates were extracted from the constructed effect size maps of each included study for the peak voxel of each of the clusters identified in each of the four meta-analysis we conducted. Using these, funnel plots were created and visually inspected. A funnel plot displays effect sizes (X-axis) against a measure of the study’s precision (i.e. sample size, Y-axis). In the absence of publication bias, studies with high precision will cluster around average effect sizes, while studies with low precision should be spread evenly on both sides of the average effect size, creating a roughly funnel-shaped distribution. Deviation from this symmetric shape can indicate publication bias (Lin and Chu, 2018) and should call for caution on interpreting such findings. We used the Egger regression test as a quantitative method of assessing asymmetry in the funnel plots. Potential publication bias is indicated if the intercept of the regression of effect size/standard error on 1/standard error significantly deviates from zero (p<0.05) (Lin and Chu, 2018).

#### 2.4.4. Meta-regressions

Previous studies have found that age (Rademacher et al., 2014) and gender (Greimel et al., 2018; Spreckelmeyer et al., 2009; Wang et al., 2017) differentially modulate the brain’s response during social and monetary incentive anticipation. For instance, one of our previous studies exploring the effects of age found that the nucleus accumbens response to cues of reward was higher for social when compared to monetary rewards in an older sample, but the opposite was true in a younger sample(Rademacher et al., 2014). Another study exploring the effects of gender showed that while in women the anticipation of monetary and social rewards engages the same brain areas, in men the anticipation of monetary rewards engaged a much wider network of brain areas in comparison to social rewards (Spreckelmeyer et al., 2009). Therefore, we considered that it was important to elucidate the impact of age and gender on the BOLD response to social reward and punishment anticipation or receipt. To achieve this, we conducted separate voxel-wise meta-regressions including mean age or the proportion of male participants across studies as moderator variables. Metaregression is a classical meta-analytic approach that allows the exploration of the impact of moderator variables on the effect sizes reported across studies using regression-based techniques (Lipsey, 2003). We took a conservative approach where we corrected the significance level for the number of meta-regressions conducted (2 metaregressions per meta-analysis), in order to contain the false positive rate. Hence, voxels with AES-SDM z-values corresponding to p-values <0.0005 (0.001/2) were considered significant, but voxels with AES-SDM z-values <1, or in clusters with less than 10 voxels, were discarded in order to reduce the false positive rate.

#### 2.4.5. Accounting for MRI signal dropout in brain areas afflicted by the susceptibility artefact

The ventromedial prefrontal cortex (vmPFC) is considered an important area for encoding social information (Hiser and Koenigs, 2018). This area is known to suffer from distortion and signal dropout during fMRI scanning due to its proximity to air and bone around the sinuses (Sutton et al., 2009). To examine whether our analyses may have missed effects due to partial coverage of this area in at least some of the studies, we examined the whole-brain coverage of the studies included in each of our main meta-analyses. This was implemented by binarising each of our effect size maps, after registration to a common template, based on whether there was signal in each voxel or not and summing these images to create coverage density maps. For those studies where we used peak coordinates, we assumed there was no signal drop out and used a whole-brain binarized template mask (note we have no way of confirming whether an area was covered, since in these cases the effect size may be null for those voxels just because they are far away from the peaks used to reconstitute the effect size maps). Since we found a decline in coverage around the anterior and inferior edge of the vmPFC in our data, we used the method developed by *Cutler and Campbell-Meiklejohn* to perform an adjusted analysis by modifying the calculations run by AES-SDM to include only studies with data present on a voxel by voxel basis (i.e. for each voxel, the adjusted meta-analysis only includes a study if it sampled that specific voxel; if no signal was present at that voxel, then this study would have not been included in the meta-analytic calculations for that specific voxel). This would allow us to check whether we might have missed significant effects in a given area simply because some of our statistical maps did not contain information in these voxels. For a detailed description of the method, please see (Cutler and Campbell-Meiklejohn, 2019).

#### 2.4.6. Conjunction analyses

We conducted two sets of separate conjunction analyses. First, in order to identify areas commonly recruited by i) the anticipation of social rewards and punishments (anticipation of social incentives), ii) the receipt of social rewards and punishment (receipt of social incentives), iii) the anticipation and receipt of social rewards (social reward processing), and iv) the anticipation and receipt of social punishments (social punishment processing), we created overlap maps between the thresholded and binarized maps of each correspondent contrasts (the resulting map represents a conjunction of all the voxels present in both meta-analytic maps). However, it should be noted that the number of studies included in the meta-analysis for each contrast was variable, and particularly unbalanced across the contrasts focusing on social reward and social punishment. It is possible that a certain voxel reaches significance in a specific meta-analytic contrast that includes a higher number of studies (e.g. such as the social reward versus neutral feedback anticipation contrast), while the same voxel does not reach significance in a different contrast simply because the number of contributing studies is smaller. While overlapping voxels can be interpreted with confidence, these differences in the number of contributing studies across contrasts preclude further inferences on areas of non-overlap or subtraction analyses.

Second, we wanted to gain insight about whether the brain areas identified in our meta-analytic maps for the social incentive delay task broadly mapped onto the brain areas identified in a similar previous meta-analysis for the monetary incentive task that has also used the AES-SDM approach (Wilson et al., 2018). Since Wilson et al. only focused on the anticipation period of the monetary incentive delay task, we created overlap maps between the meta-analytic maps for the anticipation phase of social rewards/punishment that we obtained in our study, and the meta-analytic maps for the anticipation phase for monetary rewards/punishments obtained in the previous study (Wilson et al., 2018). Similar comparisons could not be conducted for the receipt phase because there is no meta-analysis using AES-SDM in the case of rewards, and we could not retrieve the final meta-analytic maps from the authors in the case of punishment (Dugre et al., 2018b)).

#### 2.4.7. Labelling and atlases

We labelled each significant cluster using the Harvard-Oxford atlas in FSL (FMRIB Software Library, www.fmrib.ox.ac.uk/fsl).

## 2. Results

### 2.1. Included studies

Our literature search process resulted in the identification of a total of 104 studies. We identified four additional references through screening of the reference lists of the included studies. After removing duplicates, 57 studies remained. Thirty-three studies were excluded during title, abstract and key-words screening, resulting in 24 full-texts to be assessed for eligibility. From these 24 full-texts, we end up excluding six studies. Reasons for exclusion are detailed in our PRISMA flow diagram (Fig. 1). Our final pool of studies included 18 references. We contacted the lead, senior and/or corresponding authors from these 18 studies to obtain the raw statistical maps, of whom 12 responded. We excluded two additional studies for which we could not retrieve from the paper or have provided by the authors statistical maps or peak coordinates for any of the reported contrasts. Therefore, the final pool of studies that was included in our meta-analyses consisted of 16 studies.

The combined sample size across all 16 included studies was 502 participants, mean age 25.95 years old (SD 5.95), mean percentage of males 52.90% (SD 33.70), 62.5% of the studies included only right-handed participants, while for 37.5% of the studies, information on participants’ handedness was not available. Fourteen out of the 16 studies used fMRI data acquired at 3T scanners – the remainder two studies used data from 1.5T scanners. For two studies, we included data acquired after the intranasal administration of a placebo in the context of pharmacological oxytocin studies. Nine studies used outcome feedback in the form of static emotional faces, three studies in the form of dynamic videos depicting positive/negative emotional reactions to performance (happy or angry face with thumbs up or down) and four studies used verbal feedback (either auditory or written). See Table 1 for an overview of the data included in each of our four meta-analysis and table S1 for details on sociodemographics, paradigm specification, MRI data acquisition and analysis.

**Table 1.** Summary of available data for each meta-analysis. In this table, we summarize the total number of datasets used in each of our 4 meta-analyses, the number of datasets for which we could retrieve statistical parametric maps and the total pooled sample size of the datasets used in each meta-analysis.

| Contrast | N datasets | N statistical maps | Pooled sample size |
| --- | --- | --- | --- |
| 1. Anticipation of social reward | 16 | 9 | 502 |
| 2. Receipt of social reward | 9 | 6 | 272 |
| 3. Anticipation of social punishment avoidance | 5 | 3 | 129 |
| 4. Receipt of social punishment | 4 | 3 | 107 |

### 2.2. Meta-analytical findings

#### 2.2.1. Anticipation of social reward

Sixteen-studies reported results on the contrast of anticipation of social reward versus neutral feedback anticipation. We obtained statistical maps from nine of the 16 studies, and used peak coordinate statistics for the remaining seven studies. This meta-analysis included data from the full combined sample of 502 participants.

#### 2.2.2. Main meta-analytic findings

We present a detailed description of all the clusters and respective peaks corresponding to increases and decreases in the BOLD signal during the cued anticipation of social rewards versus neutral feedback in Table S2. Briefly, we found increases in the BOLD signal in a network of regions spanning the striatum, thalamus, insula and amygdala bilaterally, the right olfactory cortex, the precentral gyri and the supplementary motor area bilaterally, the left middle occipital gyrus, left inferior frontal gyrus, the brainstem, and the cerebellum (vermic lobule). We found decreases in the BOLD signal in the left median/paracingulate gyri, the right postcentral gyrus, the middle temporal gyrus bilaterally, the superior frontal gyrus bilaterally, the left middle frontal gyrus, the inferior frontal gyrus bilaterally, the left fusiform and angular gyri, the right middle occipital gyrus, the right parahippocampal gyrus, and the left cerebellum (Fig. 2).

**Fig. 2.**
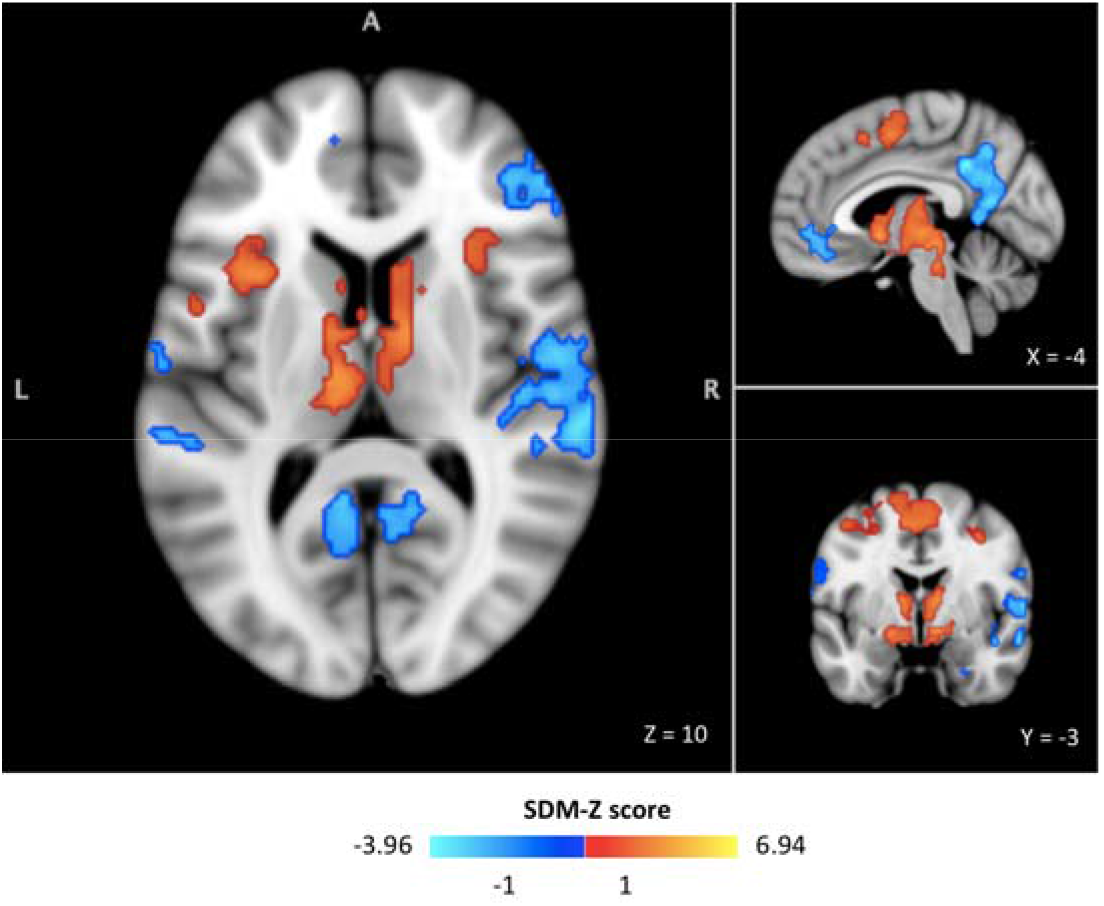
Anticipation of social reward. Meta-analytic results for the social reward anticipation contrast (16 studies, pooled sample size 502 participants). Colour bars represent SDM-Z scores. In red, we present increases and in the blue decreases in BOLD signal for the contrast anticipation of social reward versus anticipation of neutral feedback. Results were considered significant for p<0.001, SDM-Z > 1 and cluster extent > 10 voxels as per current standard recommendations for multiple comparisons control using this method.

#### 2.2.3. Sensitivity analysis

Our sensitivity analysis showed that our main meta-analytic findings are overall robust (Fig. S1).

#### 2.2.4. Heterogeneity and publication bias

We found some areas of heterogeneity in the basal ganglia, thalamus, right insula, superior temporal gyrus, midbrain and supplementary motor area (Fig. S2). We found some evidence for publication bias in five cluster peaks showing increases in the BOLD signal. These clusters included the right supplementary motor area, the left insula, the right precentral gyrus and the left middle occipital gyrus (Table S2).

#### 2.2.5. Metaregressions

We did not find any voxels where changes in BOLD were moderated by the mean age or the percentage of men in the included studies.

#### 2.2.6. Subgroup analyses

When we repeated our main meta-analysis including only studies using emotional faces as outcome (Fig. S3), or including only studies using static emotional faces (Fig. S4), or including only non-pharmacological studies (Fig. S5), we found no difference in the pattern of our results, but only observed a decrease in the size or extent of our significant clusters.

### 2.3. Receipt of social reward

Nine studies reported results on the contrast social reward feedback versus neutral feedback. We obtained statistical maps from 6 of these studies, and used peak coordinate statistics for the remaining three studies. This meta-analysis included data from 272 participants.

#### 2.3.1. Main meta-analytic findings

We present a detailed description of all clusters and respective peaks corresponding to increases and decreases in the BOLD signal during receipt of social reward versus neutral feedback in Table S3. Briefly, we found increases in the BOLD signal in a network of regions spanning the frontal medial cortex, the anterior cingulate and frontal orbital cortices bilaterally, the middle temporal gyrus bilaterally, the amygdala and hippocampus bilaterally, the right thalamus, the superior frontal gyrus bilaterally, the posterior cingulate/precuneus bilaterally, the lateral occipital cortex bilaterally, the right occipital pole and the brainstem. We found decreases in the BOLD signal in the superior frontal gyrus bilaterally, the right frontal pole, the left postcentral gyrus, the central opercular cortex/insula and the precuneus bilaterally (Fig. 3).

**Fig. 3.**
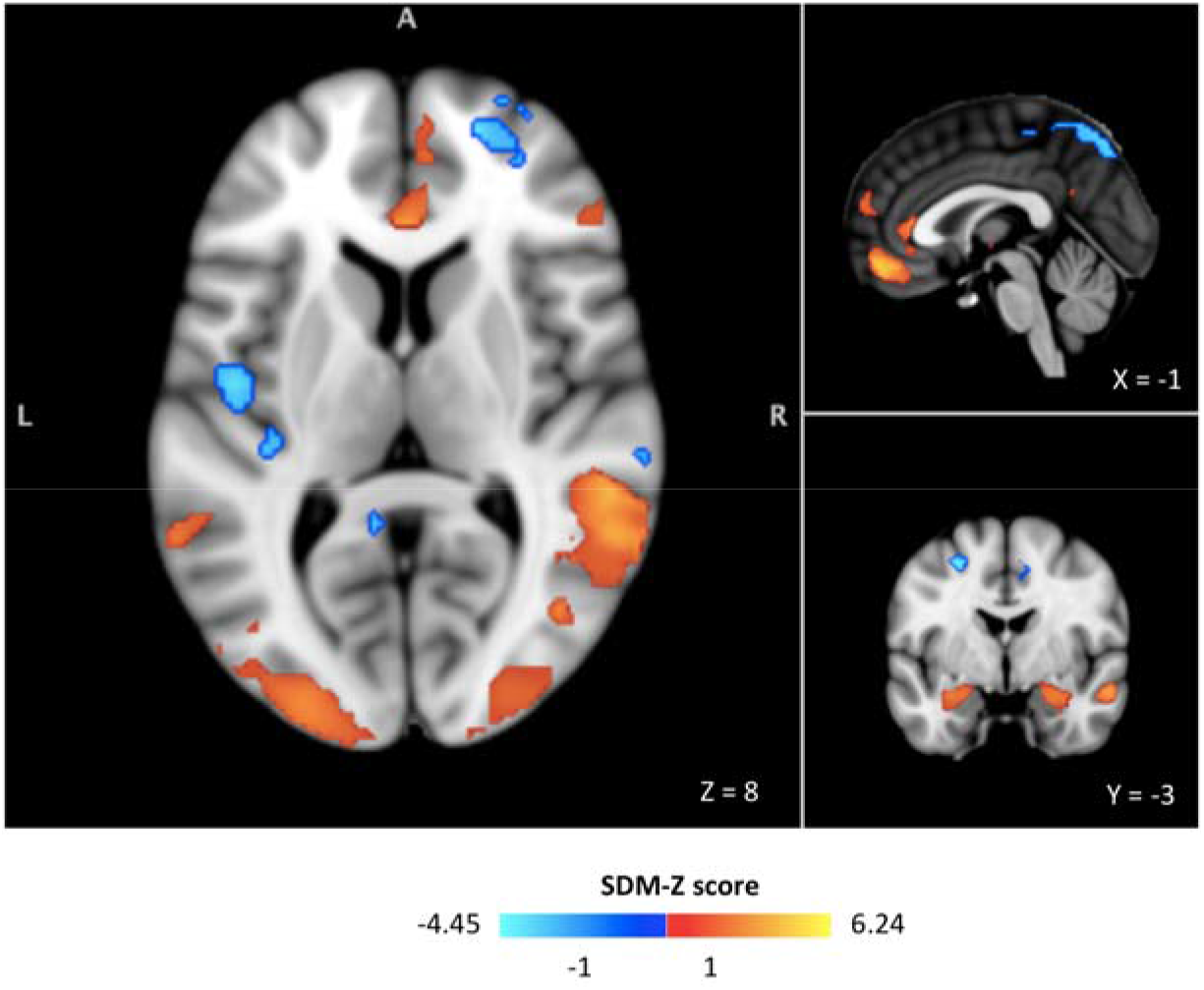
Receipt of social reward. Meta-analytic results for the social reward receipt contrast (9 studies, pooled sample size 272 participants). Colour bars represent SDM-Z scores. In red, we present increases and in the blue decreases in BOLD signal for the contrast receipt of social reward versus receipt of neutral feedback. Results were considered significant for p<0.001, SDM-Z > 1 and cluster extent > 10 voxels as per current standard recommendations for multiple comparisons control using this method.

#### 2.3.2. Sensitivity analysis

Our sensitivity analysis showed that our main meta-analytic findings were overall robust (Fig. S6).

#### 2.3.3. Heterogeneity and publication bias

We found some areas of heterogeneity mostly in the frontal medial cortex, the anterior cingulate cortex, the amygdala/hippocampus, the frontal orbital cortex, the occipital cortex and the occipital pole, and the opercular cortex/insula (Fig. S7). We found evidence for publication bias in four cluster peaks showing decreases in the BOLD signal. These clusters included the right median cingulate/paracingulate gyri, the left insula and the right inferior parietal gyri (Table S3).

#### 2.3.4. Metaregressions

We did not find any voxels where changes in the BOLD signal were moderated by the mean age or the percentage of men in the included studies.

#### 2.3.5. Subgroup analyses

When we repeated our main meta-analysis including only studies using static emotional faces (Fig. S8), or excluding studies administering placebo (Fig. S9), we found no substantial difference in the pattern of our results, other than an overall decrease in the size/extent of our significant clusters. This decrease was more pronounced when we excluded the studies administering placebo, particularly with respect to decreases in the BOLD signal.

### 2.4. Anticipation of social punishment avoidance

Only six studies reported results on the contrasts anticipation of social punishment avoidance versus neutral. We obtained statistical maps from three of these studies, and used peak coordinate statistics from two studies. This meta-analysis included data from 129 participants.

#### 2.4.1. Main meta-analytic findings

We present a detailed description of all clusters and respective peaks corresponding to increases and decreases in the BOLD signal during the anticipation of social punishment avoidance versus neutral feedback in Table S4. Briefly, we found increases in the BOLD signal in a network of regions spanning the frontal orbital cortex bilaterally, the superior and middle frontal gyri bilaterally, the amygdala, thalamus, septal nuclei, striatum and pallidum bilaterally, the right insula, the brainstem, the right lateral and fusiform occipital cortices, and the occipital pole bilaterally. We found decreases in the BOLD signal mainly in the paracingulate gyrus, the frontal pole bilaterally, the left precentral gyrus, the left temporal pole, the supramarginal gyri bilaterally, the left parietal operculum, the left middle/inferior temporal gyri and the right cerebellum (Crus I) (Fig. 4).

**Fig. 4.**
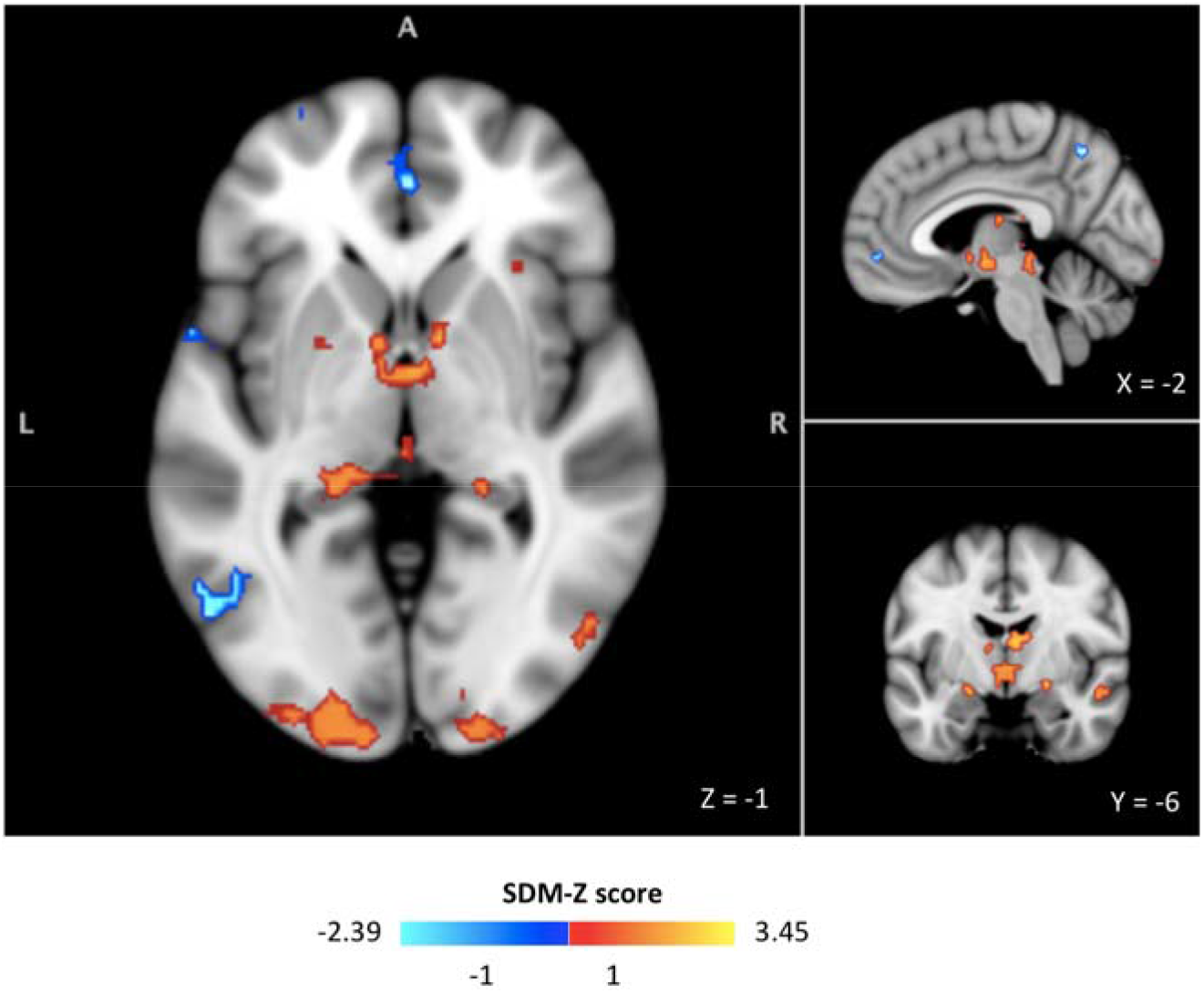
Anticipation of social punishment avoidance. Meta-analytic results for the social punishment anticipation contrast (5 studies, pooled sample size 129 participants). Colour bars represent Z-SDM scores. In red, we present increases and in the blue decreases in BOLD signal for the contrast anticipation of social punishment avoidance versus anticipation of neutral feedback. Results were considered significant for p<0.001, Z-SDM > 1 and cluster extent > 10 voxels as per current standard recommendations for multiple comparisons control using this method.

#### 2.4.2. Sensitivity analysis

Our sensitivity analysis showed that our main meta-analytic findings were overall robust (Fig. S10).

#### 2.4.3. Heterogeneity and publication bias

We found only a few areas of heterogeneity mostly in the occipital poles (bilaterally), the thalamus and the left striatum (Fig. S11). We did not find evidence for publication bias in any of the reported peaks (Table S4).

#### 2.4.4. Metaregressions

We did not find any voxels where changes in the BOLD signal were moderated by the mean age of the participants included. We found positive associations between the percentage of men included in the studies and the BOLD signal change in the occipital poles bilaterally, the right thalamus, the striatum/pallidum bilaterally, and the right frontal orbital cortex, and negative associations with the BOLD signal change in the medial frontal cortex and the left middle/inferior temporal gyri (Fig. S12).

#### 2.4.5. Sub-group analyses

When we repeated our main meta-analysis excluding the study from (Nawijn et al., 2017), we found no considerable difference in the pattern of our results, but only observed a decrease in the size or extent of our significant clusters in the basal ganglia.

### 2.5. Receipt of social punishment

Four studies reported results on the contrast social punishment feedback versus neutral feedback. We obtained statistical maps from three of these studies, and used peak coordinate statistics for one study. This meta-analysis included data from 107 participants.

#### 2.5.1. Main meta-analytic findings

We present a detailed description of all clusters and respective peaks corresponding to increases and decreases in the BOLD signal during receipt of social punishment feedback versus receipt of neutral feedback in Table S5. Briefly, we found increases in the BOLD signal in a network of regions spanning the right superior/inferior frontal gyri, the right frontal pole, the right lateral occipital cortex, the frontal orbital cortex bilaterally and the left frontal operculum/insula. We found decreases in the BOLD signal in areas spanning the precuneus, the angular gyrus, the frontal pole and the superior/middle frontal gyri bilaterally, the left caudate, and the right putamen (Fig. 5).

**Fig. 5.**
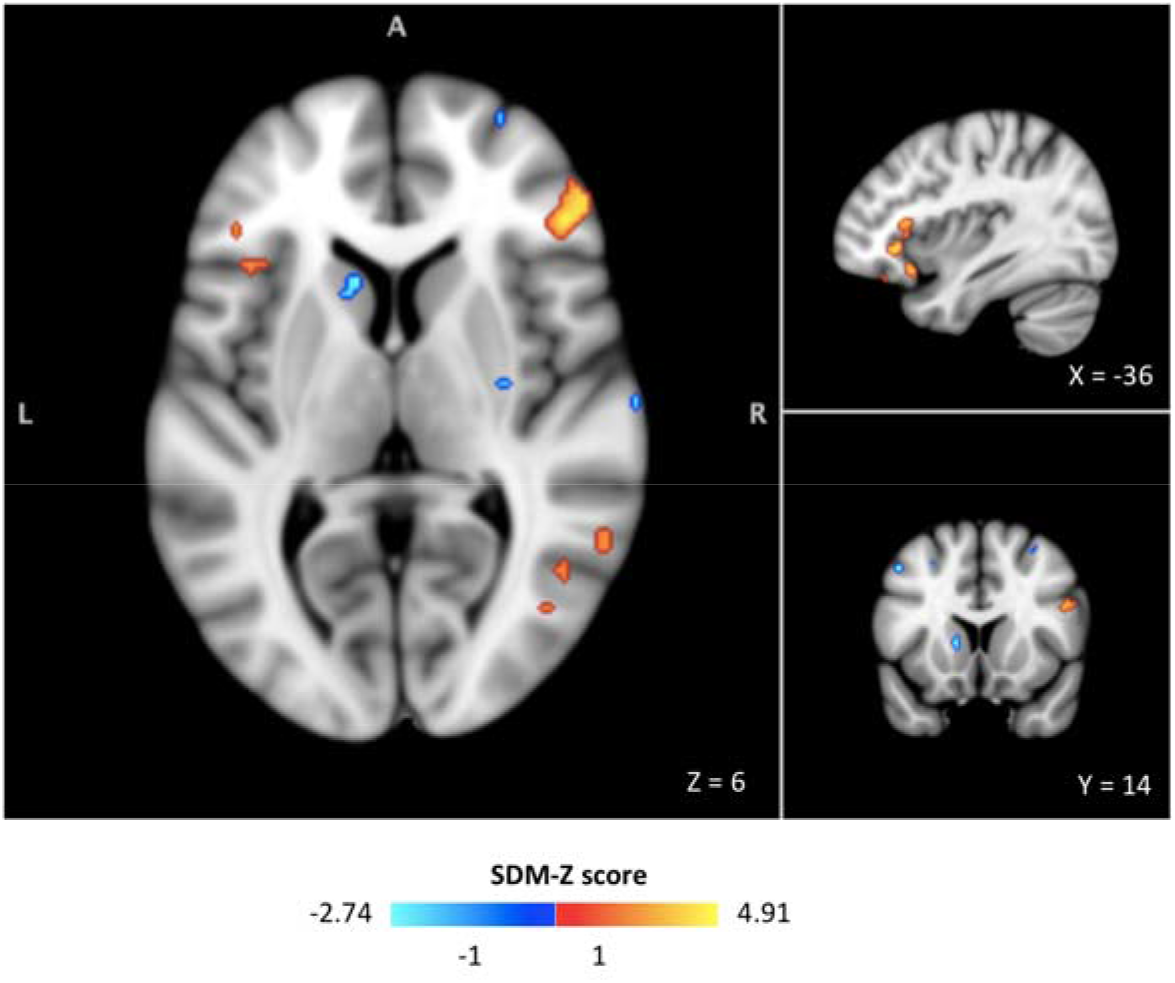
Receipt of social punishment. Meta-analytic results for the social punishment receipt contrast (4 studies, pooled sample size 107 participants). Colour bars represent Z-SDM scores. In red, we present increases and in the blue decreases in BOLD signal for the contrast receipt of social punishment versus receipt of neutral feedback. Results were considered significant for p<0.001, Z-SDM > 1 and cluster extent > 10 voxels as per current standard recommendations for multiple comparisons control using this method.

#### 2.5.2. Sensitivity analysis

Our sensitivity analysis showed that our main meta-analytic findings were overall robust for increases in the BOLD signal. However, it also showed that the robustness of the decreases in the BOLD signal was attenuated, with voxels reported in our main meta-analysis reaching significance in no more than 50% of the Jackknife leave-one-out meta-analyses (Fig. S13).

#### 2.5.3. Heterogeneity and publication bias

We found some minimal areas of heterogeneity mostly in the right caudate, the right frontal orbital cortex, the left temporal gyrus and the precuneus bilaterally (Fig. S14). We did not find evidence for publication bias in any of the reported peaks (Table S5).

#### 2.5.4. Metaregressions

We did not find any voxels where changes in the BOLD signal were moderated by mean age or the percentage of men in the included studies.

#### 2.6. Accounting for MRI signal dropout in brain areas

We did not find considerable changes in our results when we repeated our four meta-analyses accounting for MRI signal dropout in brain areas such as the ventromedial prefrontal cortex (Figs.S15 and S16). Therefore, we are unlikely to have missed increases or decreases in the BOLD signal in these areas because of signal dropout in some of the included studies.

### 2.7. Conjunction analyses to identify areas of overlap in the processing of social rewards

#### 2.7.1. Anticipation of social incentives (rewards and punishments)

The contrasts for the anticipation of social rewards and punishment avoidance (versus the anticipation of neutral feedback) overlapped in a group of brain areas that included the caudate nucleus, putamen and pallidum (bilaterally), the thalamus (bilaterally), the frontal medial cortex (bilaterally), the right insular cortex, and to a small extent the midbrain and the amygdala bilaterally (Fig. 6A).

**Fig. 6.**
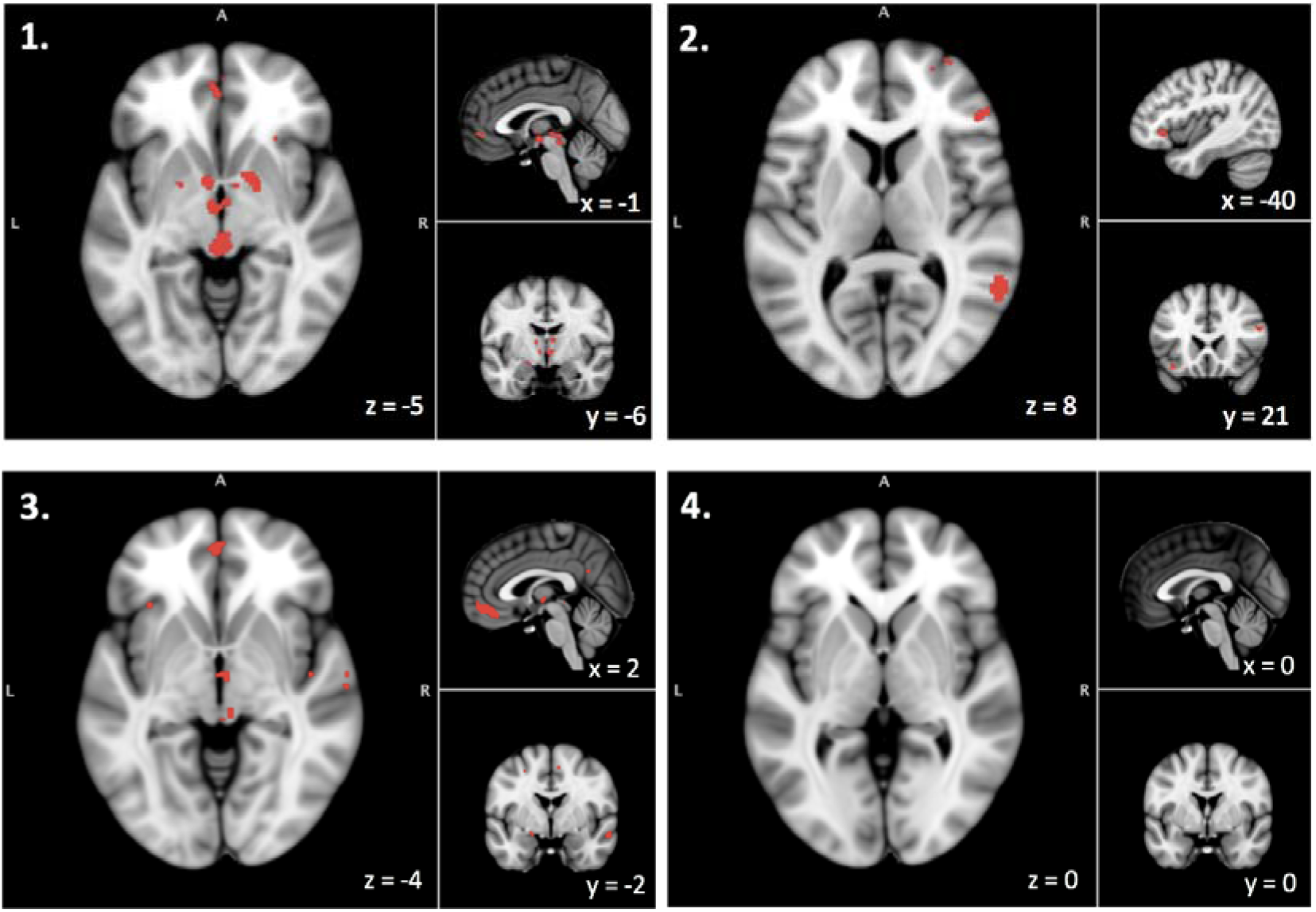
Conjunction analysis. In this figure, we present the results of a conjunction analysis examining the overlap between our four meta-analytic maps for the following pairs: 1. Anticipation of social reward AND anticipation of social punishment avoidance; 2. Receipt of social reward AND receipt of social punishment; 3. Anticipation of social reward AND receipt of social reward; 4. Anticipation of social punishment avoidance AND receipt of social punishment. Overlaps are depicted as binarized voxels coloured in red.

#### 2.7.2. Receipt of social incentives (rewards and punishments)

The contrasts for the receipt of social rewards and punishments (versus the receipt of neutral feedback) overlapped in the precuneus (bilaterally), the right lateral occipital cortex (right), and the frontal pole/orbitofrontal cortices (bilaterally) (Fig. 6B).

#### 2.7.3. Anticipation and receipt of social rewards

The contrasts for the anticipation and receipt of social rewards (versus the anticipation and receipt of neutral feedback, respectively) overlapped in the frontal medial cortex (bilaterally), the anterior (left) and middle (right) cingulate cortex, the thalamus (bilaterally), the posterior cingulate (bilaterally) and to a small extent in the left amygdala, the lateral occipital cortex (bilaterally), the superior temporal gyrus (bilaterally), the left precentral/postcentral gyri, and the right superior frontal gyrus (Fig. 6C).

#### 2.7.4. Anticipation and receipt of social punishments

We did not find any voxel where the contrasts for the anticipation and the receipt of social punishments (versus the anticipation and receipt of neutral feedback, respectively) overlapped (Fig. 6D).

### 2.8. Conjunction analyses to identify areas of overlap in the anticipation of social and monetary rewards

#### 2.8.1. Anticipation of social and monetary rewards (versus neutral feedback)

Overall, we found little evidence that the anticipation of social reward recruits modality-specific brain areas when qualitatively compared to the anticipation of monetary rewards (with the exception of the left cerebellar hemisphere – Crus I). The brain areas we identified to be involved in the anticipation of social rewards broadly map onto the same anatomical regions previously identified in (Wilson et al., 2018) to be involved in the anticipation of monetary rewards. It should be noted that in some cases the identified clusters were contiguous rather than overlapping exactly (Fig.S17).

#### 2.8.2. Anticipation of social and monetary punishments (versus neutral feedback)

Overall, we found little evidence that the anticipation of social punishment recruits modality-specific brain areas when qualitatively compared to the anticipation of monetary punishments (with the exception of the right cerebellar hemisphere – Crus I and the occipital poles). The brain areas we identified to be involved in the anticipation of social punishment avoidance broadly mapped onto the same anatomical regions identified in (Wilson et al., 2018) to be involved in the anticipation of monetary loss avoidance. It should be noted that in some cases the identified clusters were contiguous rather than overlapping exactly (Fig.S18).

## 3. Discussion

We present the results of the first comprehensive voxel-based meta-analysis using Anisotropic Effect Size Signed Differential Mapping (AES-SDM) to map the brain regions involved in the anticipation and receipt of performance dependent, probabilistic social rewards and punishments in the human brain using the SID task. All four meta-analytic maps can be freely downloaded from *Neurovault* (https://neurovault.org/collections/7793). We identify brain areas missed in individual studies due to a lack of power, including areas that are included in the default-mode network and show decreases in the BOLD signal during the anticipation of both social rewards and punishment avoidance. Additionally, we characterise the effect size and direction of changes in the BOLD signal for each brain area. Our meta-analyses enhance our understanding of the neural circuits underlying social incentive processing in healthy humans. Furthermore, our results can inform targeted hypotheses about potentially disrupted areas in the brain circuitry involved in social incentive processing that may be linked with social dysfunction in neuropsychiatric disorders. We discuss each of our main findings below.

### Brain regions involved in the anticipation of social rewards and punishment avoidance

Using a voxel-based meta-analytic method, we identified an extensive network of brain areas involved in the anticipation of social rewards. Our results consolidate findings from a previous coordinate-based meta-analysis (ALE) regarding the involvement of the basal ganglia, the midbrain, the dorsal anterior cingulate cortex, the supplementary motor area, the anterior insula and the occipital gyrus (Gu et al., 2019). However, thanks to the enhanced sensitivity of the AES-SDM meta-analytic method that we used, our results extend this network by including frontal, temporal, parietal and cerebellar regions that were not captured in the previous coordinate-based meta-analysis. Furthermore, for the first time, we identified the involvement of a robust network of brain regions typically regarded as part of the default-mode network, namely the posterior cingulate cortex, angular gyrus and inferior parietal lobe. Specifically, we found that these brain regions showed a relative decrease in the BOLD signal during the anticipation of social reward, compared to neutral feedback. These brain regions had not been previously identified in individual studies but have been suggested to participate in the brain processing of subjective value (Acikalin et al., 2017). In a similar previous AES-SDM meta-analysis of fMRI data from the MID task, the authors also identified a network of brain regions (largely overlapping with the brain region we identified here) showing decreases in the BOLD signal during the anticipation of monetary incentives (Wilson et al., 2018) that had not been identified in previous coordinate-based meta-analyses(Gu et al., 2019; Oldham et al., 2018b). Hence, our results neatly illustrate some of the advantages of the AES-SDM meta-analytic approach, including its increased sensitivity compared to the frequently underpowered single studies or coordinate-based meta-analyses (Radua and Mataix-Cols, 2012).

Furthermore, our study provides the first meta-analysis of neuroimaging data focusing on the contrast comparing the anticipation of social punishment avoidance to the anticipation on neutral feedback. We identified a robust network of brain regions showing increases and decreases in the BOLD signal similar to that involved in the anticipation of social rewards. Our conjunction analyses suggested that the anticipation of both social rewards and punishment avoidance recruits a set of well described circuits typically known as the “reward system” (Arias-Carrion et al., 2010). This observation lends support to the idea that most these areas may be part of general motivational system linked to the anticipation of highly salient incentive stimuli independent of valence or incentive-type (i.e. social or monetary). Consistent with this idea, previous findings from an ALE meta-analysis also suggested that the same network of regions are involved in the anticipation of monetary gains and losses during the MID task (Oldham et al., 2018b). However, given that our social reward and punishment avoidance anticipation meta-analyses were highly unbalanced in terms of the number of included studies, which did not allow us to conduct subtraction analyses, we should not exclude that, at least, some of the areas we identified in each of these two meta-analyses might be specifically engaged by one process or the other.

Below we discuss some of the identified brain regions that are considered to play a key role in incentive processing. We identified increases in the BOLD signal during the anticipation of social rewards and punishment avoidance in the striatum, the anterior insula, the thalamus, the anterior cingulate, the supplementary motor cortex and the amygdala. Accumulating evidence associates the ventral striatum (and the mesolimbic dopamine pathway) to motivational processing independent of stimulus valence (Brooks and Berns, 2013; Lammel et al., 2014), and many studies have suggested that the striatum plays a key role in computations that take place during social behaviour (for an extensive review please see (Baez-Mendoza and Schultz, 2013)). At the moment, it is still under debate whether the striatum solely responds to salience or might encode both salience and valence of a stimulus (Bartra et al., 2013). The anterior insula, which has dense interconnections with the striatum (Ghaziri et al., 2018), has been implicated in encoding outcome uncertainty (Gorka et al., 2016), which is a key process of the anticipation phase of the SID design. Furthermore, the insula is a key node of the salience network (Menon and Uddin, 2010) and processes ascending interoceptive and visceromotor signals (Ronchi et al., 2015). The insula is regarded as a central area involved in emotional processing and social cognition (Couto et al., 2013), including the detection of salience of social stimuli or events (Chen et al., 2009; Feng et al., 2015; Luo et al., 2018). Therefore, the involvement of the insula during the anticipation of social rewards and punishment avoidance in the SID task is compatible with an overall role of the insula in the autonomic activation (arousal) associated with the anticipation of salient social incentives (Schneider et al., 2018). The thalamus encodes an alerting signal to respond to salient stimuli (Wolff and Vann, 2019; Zhu et al., 2018) by conveying interoceptive information from the insula to the striatum, where an appropriate action response is then selected (Huang et al., 2018). Interestingly, social cognition impairments have been reported in stroke patients with unilateral thalamic lesions (Wilkos et al., 2015). Given that the SID is a motor response time task, the presence of the supplementary motor cortex, typically recruited during movement planning and control (Tanji, 1994), is consistent with the characteristics of the task. Additionally, recent studies have shown that the motor and premotor cortices also encode reward signals related to both the anticipation and receipt of a reward (Ramakrishnan et al., 2017; Ramkumar et al., 2016). The dorsal cingulate is a region that subserves both cognition and motor control (Beckmann et al., 2009) and a plethora of studies have implicated this area in processes such as attention for action/target selection (Hayden and Platt, 2010; Isomura et al., 2003), motivation (Monosov, 2017), motor response selection (Badgaiyan and Posner, 1998), performance monitoring (Gehring and Knight, 2000), novelty detection (Hayden et al., 2011) and social cognition (namely, in tracking others’ motivation) (Apps et al., 2016). Lastly, the amygdala, which has been classically linked with negative emotional processing, has more recently been proposed to respond to stimulus salience/arousal rather than valence (Bonnet et al., 2015; Fastenrath et al., 2014; Zheng et al., 2017). In addition, the amygdala has also been proposed to be instrumental during the processing of emotional and socially relevant information (including the processing of emotions from faces (Todorov, 2012b)).

For the first time, we also identified a group of brain regions showing decreases in the BOLD signal during the anticipation of both social rewards and punishment avoidance. These regions are part of what is commonly described as the default-mode network. There has been debate about the exact functions of the default-mode network (Crittenden et al., 2015; Raichle, 2015). Current views subdivide this network into a core subsystem, anchored in the posterior cingulate cortex and the anteromedial prefrontal cortex, which has been associated with self-referential processes; a medial temporal subsystem related to the processing of past and future autobiographical thoughts; and a dorsal medial subsystem anchored in the dorsomedial PFC, related to social cognition, story comprehension and semantic processing (Salomon et al., 2014). Importantly, the default mode network has been associated with decreases in the BOLD signal during external-oriented active tasks (Anticevic et al., 2012), thus leading some researchers to label the network as a task-negative network (this conceptualization of the default-mode network as a task-negative network has nevertheless been recently contested - see (Spreng, 2012)). Given that the SID task is an active task, the pattern of decreases in the BOLD signal that we observed is consistent with the idea that the brain should disengage from self-oriented processing to attend to relevant external stimuli (Scheibner et al., 2017). This idea is further supported by the activation of key nodes of the salience network such as the insula and the anterior cingulate cortex that we reported. These nodes are typically assumed to participate in the switch between self- and external-oriented processing (Corbetta and Shulman, 2002).

One key question in the field regards the extent to which the processing of social incentives involves additional specific brain regions (Gu et al., 2019; Izuma et al., 2008) that are typically not engaged during the processing of other types of incentives (i.e. monetary). One prominent hypothesis states that the anticipation and receipt of social incentives engages both a generalist neural network that consists of brain regions involved in the processing of incentives irrespective of their type (such as the basal ganglia), and a specialist network of regions specifically involved in the processing of social information (such as the temporoparietal junction, the dorsomedial prefrontal cortex, the precuneus, and the superior temporal gyrus) (Barman et al., 2015a; Goerlich et al., 2017a; Spreckelmeyer et al., 2013a). This hypothesis has been challenged though by at least two recent meta-analyses, the results of which were consistent with the idea that a general-purpose brain system is involved in the anticipation of incentives irrespective of their type. Specifically, the first meta-analysis showed that the circuits engaged during the anticipation of monetary gains also encompass some of the brain areas often attributed to social information processing(Wilson et al., 2018). The second meta-analysis compared the anticipation of monetary and social rewards and failed to find any significant differences in the underpinning brain regions (Gu et al., 2019).

To help illuminate this question, we conducted an exploratory conjunction analysis between the meta-analytic effect size maps on the SID task from our study and meta-analytic effect size maps on the MID task that we obtained from another study that used the same AES/SDM meta-analytic method (Wilson et al., 2018). We only focused on the anticipation period as the previous study had not analysed data from the outcome phase of the MID task. Overall, we found little evidence of modality-specific processing, supporting the idea that, at least regarding the anticipation of incentives, a general-purpose anticipation system is recruited by both monetary and social incentives. However, in some cases we noted the recruitment of voxels across tasks that may have belonged to the same brain regions but were anatomically contiguous rather than overlapping. It is therefore tempting to speculate that rather than recruiting different neural systems, the anticipation of social and monetary incentives may be functionally encoded in the same regions but engage different neuronal ensembles. Indeed, something similar has been demonstrated for the brain encoding of physical and social rejection, which are characterized by differential multivariate voxel patterns despite common fMRI activity at the gross anatomical level (Woo et al., 2014). This is a hypothesis that should be addressed in future studies combining the application of both tasks in the same individuals with current state-of-art multivoxel pattern recognition techniques to ascertain whether modality can be decoded by a classifier trained on brain voxel-based responses (Corradi-Dell’Acqua et al., 2016; Mahmoudi et al., 2012).

### Brain regions engaged during the receipt of social rewards and punishments

Our study provides the first meta-analysis of neuroimaging data focusing on the contrasts comparing the receipt of social rewards or punishments to neutral feedback. We identified a robust network of brain regions engaged during the receipt of social rewards, including the ventromedial frontal (vmPFC) and orbitofrontal cortices, the anterior cingulate cortex, the amygdala, the hippocampus, the occipital cortex and the brainstem. Below we discuss these findings in detail. Increases in BOLD signal in the ventromedial prefrontal cortex and the orbitofrontal cortex during the receipt of social rewards have also been consistently associated with the receipt of monetary outcomes (Oldham et al., 2018b). Previous work has suggested that the vmPFC and the orbitofrontal cortex encode subjective value (Piva et al., 2019) and the associative relationship between stimuli and outcome (de Wit et al., 2009). Furthermore, increasing evidence has implicated the vmPFC in multiple aspects of social cognition, such as facial emotion recognition, theory-of-mind ability, and the processing self-relevant information (for an extensive review and meta-analysis see (Hiser and Koenigs, 2018)).

Furthermore, we also observed increases in the BOLD signal in the amygdala, the brainstem, the anterior cingulate cortex, and the occipital cortex. The amygdala has been proposed to participate in the representation of the utility or affective value during monetary incentives receipt(Ernst et al., 2005; Hampton et al., 2007). According to this model, the utility encoded by the amygdala is thought to be then used to inform behavioural and physiological responses towards and away from positive and negative stimuli, respectively. This process has been suggested to include dispatches of the information encoding the affective value of stimuli from the amygdala to different brain systems, including the arousal circuits of the brainstem (Ernst et al., 2005; Hampton et al., 2007). Increased BOLD signal in the amygdala, the anterior cingulate cortex, and the occipital cortex is also a consistent finding of studies investigating brain responses to emotional faces(Fusar-Poli et al., 2009; Todorov, 2012a), which were the most frequently used outcome stimuli in the receipt phase of the SID tasks we included in our meta-analyses.

One notable absence was the lack of evidence for the involvement of the basal ganglia during the receipt of social rewards. In contrast, previous meta-analytic evidence has shown that the ventral striatum is engaged during the receipt of monetary rewards in the MID task (Oldham et al., 2018b). Even though the MID and the SID tasks are not classical learning tasks, the ventral striatal response to uncertain monetary rewards has been commonly interpreted in the reinforcement learning literature as the neural correlate of the dopaminergic neurons coding a prediction error signal (Nasser et al., 2017). Similar prediction errors have been reported to occur in the striatum during social learning (Joiner et al., 2017). Therefore, we would have expected to have found an increase in the BOLD signal in the ventral striatum during the receipt of social rewards. Indeed, an increase in the BOLD signal in the ventral striatum was noted in some of the leave-one-out meta-analyses that were part of the sensitivity analysis; however, this increase was not consistent enough to be captured in the main meta-analysis. Ultimately, this could have resulted from the fact that most studies used high success rates (~66.6%). These high success rates might have biased the expectations of participants about the outcomes towards high levels of certainty. According to the prediction error theory presented above, if participants were expecting to be rewarded with high levels of certainty and the outcome matched participants’ expectations about reward outcome most of the times, then one would expect the BOLD response in the basal ganglia to be reduced. While the same principle would apply for the MID task, we note that the number of studies included in previous MID meta-analyses was considerably higher than those of the studies using its SID counterpart we included here. Hence, it is possible we might have not had enough power to detect smaller changes in the BOLD signal in the basal ganglia during the receipt of social rewards in our meta-analysis.

Regarding the brain regions engaged during the receipt of social punishments, we identified a network of brain regions showing increases in the BOLD signal, including the orbitofrontal cortex, the superior and inferior frontal gyri, the lateral occipital cortex and the insula. Additionally, and in contrast to the receipt of social rewards, we also observed a decrease in the BOLD signal in the basal ganglia in response to the receipt of social punishments (compared to neutral feedback). This finding is in line with previous evidence that monetary loss, as compared to neutral feedback, results in a decrease in the BOLD signal in the striatum (in contrast to monetary gain, which results in an increase in the BOLD signal) (Delgado et al., 2000). The sensitivity analyses indicated that we should consider the meta-analytic results from this contrast with caution, as they indicated lack of robustness (that is, the reported findings could be driven by the specific combination of studies). This is not unexpected given the small number of studies (N=4) included in the meta-analysis for this contrast.

Our conjunction analysis identified clusters that were commonly engaged during the receipt of both social rewards and punishments. These clusters extended over the lateral occipital cortex, the precuneus, the cuneal cortex, the frontal pole and the orbitofrontal cortex. This observation is compatible with the hypothesis that certain regions may be engaged in shared sensory/cognitive processes irrespective of the valence of the feedback, and in encoding the subjective relevance of the feedback (Oldham et al., 2018b).

### Publication bias and heterogeneity

Very few neuroimaging meta-analyses consider the issues of publication bias and heterogeneity, which are commonly addressed in behavioural meta-analyses (Muller et al., 2018; Radua and Mataix-Cols, 2012). The consideration of publication bias in neuroimaging meta-analysis such as the ones we present here calls for reflection on one main issue. Voxels whose effect may have failed to survive multiple comparisons in individual studies would have been assigned an estimated effect size of zero when only peak coordinates are available, compared to approaches using whole brain statistical maps. With this in mind, we conducted and present the results of a publication bias analysis using both the inspection of funnel plots and the Egger’s test on the effects extracted for each peak of our four meta-analyses. We could only find some indication for publication bias for some of the peaks identified in the meta-analysis of the anticipation of social reward versus anticipation of neutral feedback contrast. For these peaks, the inspection of the respective funnel plots indicated the lack of smaller studies reporting small effect sizes for these peaks. Therefore, the findings regarding these peaks should be taken with caution. However, we should note that none of these peaks would have shown publication bias if we had applied statistical correction in our publication bias analyses for the number of peaks examined. Regarding the issue of methodological heterogeneity across the studies included in the reported meta-analyses, we found evidence suggesting the existence of considerable methodological heterogeneity in the meta-analytic maps for the anticipation of social reward or punishment avoidance versus the anticipation of neutral feedback contrast, and the receipt of social rewards versus neutral feedback contrasts. For the anticipation of social punishment avoidance versus the anticipation of neutral feedback contrast, this heterogeneity is likely to be explained by differences in the percentage of men included across studies, as suggested by our exploratory metaregression analyses using the percentage of men included in each study as predictor. This finding is largely in line with previous studies showing gender-related differences in the neurophysiological underpinnings of social incentive processing (Greimel et al., 2018; Spreckelmeyer et al., 2009; Wang et al., 2017). For instance, one study in adolescents showed that during the anticipation of potential social punishment, adolescent boys, compared with girls, exhibited a reduced stimulus-preceding negativity (Greimel et al., 2018). For the remaining contrasts, we found that neither differences in the mean age nor in the percentage of men could explain the observed heterogeneity. However, it should be noted that the inclusion of both statistical maps and coordinate-based data in AES-SDM may inflate heterogeneity estimates (Radua et al., 2012).

### Limitations

Our study has some limitations we should acknowledge. First, we note the relatively small number of studies investigating the processing of social punishment using the SID task. This aspect limits the power of our meta-analyses for the anticipation and receipt of social punishment contrasts. Nevertheless, we decided to proceed with the meta-analyses of these contrasts since we were able to retrieve statistical maps for more than half of the eligible studies. In fact, the pooled sample size of our social punishment meta-analyses exceeded 100 individuals, making it the largest dataset to date investigating this question. Given the role that differences in the processing of social punishments may play in a wide range of neuropsychiatric disorders (such as major depressive disorder (Kumar et al., 2017) or antisocial personality disorder (Gong et al., 2019)), it seems imperative that future studies start to invest more in investigating the neural underpinnings of social punishment processing in both healthy and clinical samples. Second, our findings are limited by the reported sample demographics. Across all included studies, we were only able to systematically retrieve information on age and gender. While we did explore the impact of mean age and percentage of men using meta-regression, other potentially interesting variables such as substance use, education, intelligence quotient, socioeconomic status and ethnicity could not be consistently retrieved to allow for a thorough investigation of their impact. Third, our results may not generalize to the processes involved in the anticipation or receipt of social rewards or social punishments in contexts outside of the SID task, e.g. in neuroeconomic games involving the exchange of social rewards and punishments between individuals (such as the Prisoner’s dilemma (Sun et al., 2016)), or other tasks where feedback is not performance dependent (Hsu et al., 2018). Fourth, the cut-off date the studies that were included in our meta-analyses were identified was about 2 years ago, which raises the question of whether our meta-analyses might be missing a considerable number of fMRI studies using the SID task published in the past 2 years. To address this concern, we conducted a new literature search on 17/04/2020 which identified only a single study meeting our inclusion/exclusion criteria (He et al., 2019). We are thus confident that our meta-analyses still represent the vast majority of the published studies in this field.

Finally, we note that there was considerable heterogeneity across the various implementations of the SID task. For example, studies varied in terms of using static faces, dynamic faces or verbal feedback as the social stimulus, in terms of using scrambled faces, dysmorphed faces, or simple win/no-win symbols or landscape images in the neutral feedback comparison condition. This heterogeneity can be both a blessing and a curse. The use of a range of stimuli is more representative of the richness of real-life social communication and enhances our confidence that the reported results are not confined to specific lab parameters. At the same, this heterogeneity renders meta-analytic efforts more difficult as the impact of relevant task variations must be examined, but this cannot be accomplished until a sufficient number of studies has accumulated. One aspect of task heterogeneity that we would like to highlight regards the probabilistic structure of the task. Like the MID task, the SID task is performance dependent. An adjusting algorithm alters the response window so that it maintains a relatively sTable Success rate across participants (typically 67% in the MID task). Similarly, in 4 out of 5 studies that included a social punishment condition, the success rate ranged between 60-67%, suggesting that social punishment *avoidance* is the predominant anticipated outcome in this condition, which may be interpreted as an inherently rewarding outcome, that is, as an instance of negative reinforcement (Kim et al., 2006). In one study though(Nawijn et al., 2017), the success rate was set to 34%, rendering social punishment *in itself* is the predominant anticipated outcome in this condition. This may result in a range of different neurocognitive processes experienced during anticipation and outcome presentation, including emotional responses (e.g. frustration), perception of self-efficacy, and even feelings of acquired learned helplessness driven by the enduring repeated experience of punishment that might be perceived to be beyond the participants’ own control (Wanke and Schwabe, 2019). We believe that future studies need to systematically investigate the impact of this aspect of heterogeneity at the neural level, as understanding the brain circuitry involved in the processing of negative reinforcers and punishments (as opposed to rewards) may be particularly useful for a range of neuropsychiatric disorders, such as depression.

## 4. Conclusion

This is the first voxel-based meta-analysis mapping the brain regions involved in the anticipation and receipt of social rewards and punishments in the human brain, as captured by the SID task. We identify brain areas missed in individual studies due to a lack of power, such as decreases in the BOLD signal in areas that are part of the default-mode network during the anticipation of both social rewards and punishment avoidance. We also characterise the effect size and direction of changes in the BOLD signal for each brain area. Qualitative anatomical comparisons showed little evidence supporting the involvement of domain-specific brain areas during the anticipation of social incentives, lending support to the hypothesis that a shared neural circuit underpins the anticipation of incentives irrespective of their type (i.e. social versus monetary). We noted the scarcity of studies focusing on the processing of social punishments despite the importance of this condition for several neuropsychiatric disorders. Our results provide a stereotaxic set of brain regions which could serve as regions-of-interest for future hypothesis-driven research seeking to investigate further how the human brain processes social incentives and how the disruption of these processes might contribute to the social dysfunction observed across many neuropsychiatric disorders. Ultimately, this knowledge may help us to identify potential target circuits that may be modulated by therapeutic interventions aiming to restore dysfunctional social incentive processing during disease.

## Author contributions

Conceptualization: DM, ASG and YP; Data collection: DM, LR, RT, JAR, DS, KSG, LN, HRC, RW and SB; Data analysis: DM; Writing of first draft: DM and YP; Revision of the final manuscript: all authors revised the final version of the manuscript for intellectual content and approved the submitted version.

## Acknowledgments

We would like to thank Miss Fizza Qureshi for her help during data retrieval and Miss Jo Cutler for sharing her code to implement AES-SDM meta-analysis adjusting for dropout. This study was part-funded by an Economic and Social Research Council Grant (ES/K009400/1) to YP. This manuscript represents independent research.

## References

Acikalin, M.Y., Gorgolewski, K.J., Poldrack, R.A., 2017. A Coordinate-Based Meta-Analysis of Overlaps in Regional Specialization and Functional Connectivity across Subjective Value and Default Mode Networks. Front Neurosci 11, 1.

Anticevic, A., Cole, M.W., Murray, J.D., Corlett, P.R., Wang, X.J., Krystal, J.H., 2012. The role of default network deactivation in cognition and disease. Trends in Cognitive Sciences 16, 584–592.

Apps, M.A., Rushworth, M.F., Chang, S.W., 2016. The Anterior Cingulate Gyrus and Social Cognition: Tracking the Motivation of Others. Neuron 90, 692–707.

Arias-Carrion, O., Stamelou, M., Murillo-Rodriguez, E., Menendez-Gonzalez, M., Poppel, E., 2010. Dopaminergic reward system: a short integrative review. Int Arch Med 3, 24.

Badgaiyan, R.D., Posner, M.I., 1998. Mapping the cingulate cortex in response selection and monitoring. Neuroimage 7, 255–260.

Baez-Mendoza, R., Schultz, W., 2013. The role of the striatum in social behavior. Frontiers in Neuroscience 7.

Barman, A., Richter, S., Soch, J., Deibele, A., Richter, A., Assmann, A., Wustenberg, T., Walter, H., Seidenbecher, C.I., Schott, B.H., 2015a. Gender-specific modulation of neural mechanisms underlying social reward processing by Autism Quotient. Social Cognitive and Affective Neuroscience 10, 1537–1547.

Barman, A., Richter, S., Soch, J., Deibele, A., Richter, A., Assmann, A., Wustenberg, T., Walter, H., Seidenbecher, C.I., Schott, B.H., 2015b. Gender-specific modulation of neural mechanisms underlying social reward processing by Autism Quotient. Soc Cogn Affect Neurosci 10, 1537–1547.

Bartra, O., McGuire, J.T., Kable, J.W., 2013. The valuation system: a coordinate-based meta-analysis of BOLD fMRI experiments examining neural correlates of subjective value. Neuroimage 76, 412–427.

Beckmann, M., Johansen-Berg, H., Rushworth, M.F.S., 2009. Connectivity-Based Parcellation of Human Cingulate Cortex and Its Relation to Functional Specialization. Journal of Neuroscience 29, 1175–1190.

Bonnet, L., Comte, A., Tatu, L., Millot, J.L., Moulin, T., de Bustos, E.M., 2015. The role of the amygdala in the perception of positive emotions: an “intensity detector”. Frontiers in Behavioral Neuroscience 9.

Brooks, A.M., Berns, G.S., 2013. Aversive stimuli and loss in the mesocorticolimbic dopamine system. Trends in Cognitive Sciences 17, 281–286.

Button, K.S., Ioannidis, J.P., Mokrysz, C., Nosek, B.A., Flint, J., Robinson, E.S., Munafo, M.R., 2013. Power failure: why small sample size undermines the reliability of neuroscience. Nature reviews. Neuroscience 14, 365–376.

Chen, Y.H., Dammers, J., Boers, F., Leiberg, S., Edgar, J.C., Roberts, T.P.L., Mathiak, K., 2009. The temporal dynamics of insula activity to disgust and happy facial expressions: A magnetoencephalography study. Neuroimage 47, 1921–1928.

Corbetta, M., Shulman, G.L., 2002. Control of goal-directed and stimulus-driven attention in the brain. Nature Reviews Neuroscience 3, 201–215.

Corradi-Dell’Acqua, C., Tusche, A., Vuilleumier, P., Singer, T., 2016. Cross-modal representations of first-hand and vicarious pain, disgust and fairness in insular and cingulate cortex. Nature Communications 7.

Couto, B., Sedeno, L., Sposato, L.A., Sigman, M., Riccio, P.M., Salles, A., Lopez, V., Schroeder, J., Manes, F., Ibanez, A., 2013. Insular networks for emotional processing and social cognition: Comparison of two case reports with either cortical or subcortical involvement. Cortex; a journal devoted to the study of the nervous system and behavior 49, 1420–1434.

Cremers, H.R., Veer, I.M., Spinhoven, P., Rombouts, S.A.R.B., Roeiofs, K., 2015. Neural sensitivity to social reward and punishment anticipation in social anxiety disorder. Frontiers in Behavioral Neuroscience 8.

Cremers, H.R., Wager, T.D., Yarkoni, T., 2017. The relation between statistical power and inference in fMRI. Plos One 12.

Crittenden, B., Mitchell, D.J., Duncan, J., 2015. Recruitment of the default mode network during a demanding act of executive control. Elife 4.

Cutler, J., Campbell-Meiklejohn, D., 2019. A comparative fMRI meta-analysis of altruistic and strategic decisions to give. Neuroimage 184, 227–241.

de Wit, S., Corlett, P.R., Aitken, M.R., Dickinson, A., Fletcher, P.C., 2009. Differential Engagement of the Ventromedial Prefrontal Cortex by Goal-Directed and Habitual Behavior toward Food Pictures in Humans. Journal of Neuroscience 29, 11330–11338.

Delgado, M.R., Nystrom, L.E., Fissell, C., Noll, D.C., Fiez, J.A., 2000. Tracking the hemodynamic responses to reward and punishment in the striatum. J Neurophysiol 84, 3072–3077.

Delmonte, S., Balsters, J.H., McGrath, J., Fitzgerald, J., Brennan, S., Fagan, A.J., Gallagher, L., 2012. Social and monetary reward processing in autism spectrum disorders. Molecular Autism 3.

Dugre, J.R., Dumais, A., Bitar, N., Potvin, S., 2018a. Loss anticipation and outcome during the Monetary Incentive Delay Task: a neuroimaging systematic review and meta-analysis. PeerJ 6, e4749.

Dugre, J.R., Dumais, A., Bitar, N., Potvin, S., 2018b. Loss anticipation and outcome during the Monetary Incentive Delay Task: a neuroimaging systematic review and meta-analysis. Peerj 6.

Ernst, M., Nelson, E.E., Jazbec, S., McClure, E.B., Monk, C.S., Leibenluft, E., Blair, J., Pine, D.S., 2005. Amygdala and nucleus accumbens in responses to receipt and omission of gains in adults and adolescents. Neuroimage 25, 1279–1291.

Fastenrath, M., Coynel, D., Spalek, K., Milnik, A., Gschwind, L., Roozendaal, B., Papassotiropoulos, A., de Quervain, D.J.F., 2014. Dynamic Modulation of Amygdala-Hippocampal Connectivity by Emotional Arousal. Journal of Neuroscience 34, 13935–13947.

Fehr, E., Camerer, C.F., 2007. Social neuroeconomics: the neural circuitry of social preferences. Trends Cogn Sci 11, 419–427.

Feng, C.L., Luo, Y.J., Krueger, F., 2015. Neural Signatures of Fairness-Related Normative Decision Making in the Ultimatum Game: A Coordinate-Based Meta-Analysis. Human Brain Mapping 36, 591–602.

Fusar-Poli, P., Placentino, A., Carletti, F., Landi, P., Allen, P., Surguladze, S., Benedetti, F., Abbamonte, M., Gasparotti, R., Barale, F., Perez, J., McGuire, P., Politi, P., 2009. Functional atlas of emotional faces processing: a voxel-based meta-analysis of 105 functional magnetic resonance imaging studies. J Psychiatry Neurosci 34, 418–432.

Gehring, W.J., Knight, R.T., 2000. Prefrontal-cingulate interactions in action monitoring. Nature Neuroscience 3, 516–520.

Ghaziri, J., Tucholka, A., Girard, G., Boucher, O., Houde, J.C., Descoteaux, M., Obaid, S., Gilbert, G., Rouleau, I., Nguyen, D.K., 2018. Subcortical structural connectivity of insular subregions. Scientific Reports 8.

Goerlich, K.S., Votinov, M., Lammertz, S.E., Winkler, L., Spreckelmeyer, K.N., Habel, U., Grunder, G., Gossen, A., 2017a. Effects of alexithymia and empathy on the neural processing of social and monetary rewards. Brain Structure & Function 222, 2235–2250.

Goerlich, K.S., Votinov, M., Lammertz, S.E., Winkler, L., Spreckelmeyer, K.N., Habel, U., Grunder, G., Gossen, A., 2017b. Effects of alexithymia and empathy on the neural processing of social and monetary rewards. Brain Struct Funct 222, 2235–2250.

Gong, X., Brazil, I.A., Chang, L.J., Sanfey, A.G., 2019. Psychopathic traits are related to diminished guilt aversion and reduced trustworthiness during social decision-making. Scientific Reports 9.

Gorka, S.M., Nelson, B.D., Phan, K.L., Shankman, S.A., 2016. Intolerance of uncertainty and insula activation during uncertain reward. Cognitive Affective & Behavioral Neuroscience 16, 929–939.

Greimel, E., Bakos, S., Landes, I., Tollner, T., Bartling, J., Kohls, G., Schulte-Korne, G., 2018. Sex differences in the neural underpinnings of social and monetary incentive processing during adolescence. Cognitive, affective & behavioral neuroscience 18, 296–312.

Gu, R.L., Huang, W.H., Camilleri, J., Xu, P.F., Wei, P., Eickhoff, S.B., Feng, C.L., 2019. Love is analogous to money in human brain: Coordinate-based and functional connectivity meta-analyses of social and monetary reward anticipation. Neuroscience and Biobehavioral Reviews 100, 108–128.

Hampton, A.N., Adolphs, R., Tyszka, M.J., O’Doherty, J.P., 2007. Contributions of the amygdala to reward expectancy and choice signals in human prefrontal cortex. Neuron 55, 545–555.

Hayden, B.Y., Heilbronner, S.R., Pearson, J.M., Platt, M.L., 2011. Surprise Signals in Anterior Cingulate Cortex: Neuronal Encoding of Unsigned Reward Prediction Errors Driving Adjustment in Behavior. Journal of Neuroscience 31, 4178–4187.

Hayden, B.Y., Platt, M.L., 2010. Neurons in Anterior Cingulate Cortex Multiplex Information about Reward and Action. Journal of Neuroscience 30, 3339–3346.

He, Z., Zhang, D., Muhlert, N., Elliott, R., 2019. Neural substrates for anticipation and consumption of social and monetary incentives in depression. Soc Cogn Affect Neurosci 14, 815–826.

Higgins, J.P., Thompson, S.G., Deeks, J.J., Altman, D.G., 2003. Measuring inconsistency in meta-analyses. BMJ (Clinical research ed.) 327, 557–560.

Hiser, J., Koenigs, M., 2018. The Multifaceted Role of the Ventromedial Prefrontal Cortex in Emotion, Decision Making, Social Cognition, and Psychopathology. Biol Psychiatry 83, 638–647.

Hsu, C.T., Sims, T., Chakrabarti, B., 2018. How mimicry influences the neural correlates of reward: An fMRI study. Neuropsychologia 116, 61–67.

Huang, A.S., Mitchell, J.A., Haber, S.N., Alia-Klein, N., Goldstein, R.Z., 2018. The thalamus in drug addiction: from rodents to humans. Philosophical Transactions of the Royal Society B-Biological Sciences 373.

Isomura, Y., Ito, Y., Akazawa, T., Nambu, A., Takada, M., 2003. Neural coding of “attention for action” and “response selection” in primate anterior cingulate cortex. Journal of Neuroscience 23, 8002–8012.

Izuma, K., Saito, D.N., Sadato, N., 2008. Processing of social and monetary rewards in the human striatum. Neuron 58, 284–294.

Joiner, J., Piva, M., Turrin, C., Chang, S.W.C., 2017. Social learning through prediction error in the brain. NPJ Sci Learn 2, 8.

Joober, R., Schmitz, N., Annable, L., Boksa, P., 2012. Publication bias: What are the challenges and can they be overcome? Journal of Psychiatry & Neuroscience 37, 149–152.

Kim, H., Shimojo, S., O’Doherty, J.P., 2006. Is avoiding an aversive outcome rewarding? Neural substrates of avoidance learning in the human brain. Plos Biology 4, 1453–1461.

Kim, H.Y., 2015. Statistical notes for clinical researchers: Type I and type II errors in statistical decision. Restor Dent Endod 40, 249–252.

Kohls, G., Antezana, L., Mosner, M.G., Schultz, R.T., Yerys, B.E., 2018. Altered reward system reactivity for personalized circumscribed interests in autism. Molecular Autism 9.

Kohls, G., Schulte-Ruther, M., Nehrkorn, B., Muller, K., Fink, G.R., Kamp-Becker, I., Herpertz-Dahlmann, B., Schultz, R.T., Konrad, K., 2013. Reward system dysfunction in autism spectrum disorders. Soc Cogn Affect Neurosci 8, 565–572.

Kumar, P., Waiter, G.D., Dubois, M., Milders, M., Reid, I., Steele, J.D., 2017. Increased neural response to social rejection in major depression. Depression and Anxiety 34, 1049–1056.

Lammel, S., Lim, B.K., Malenka, R.C., 2014. Reward and aversion in a heterogeneous midbrain dopamine system. Neuropharmacology 76, 351–359.

Li, X., Li, Z., Li, K., Zeng, Y.W., Shi, H.S., Xie, W.L., Yang, Z.Y., Lui, S.S.Y., Cheung, E.F.C., Leung, A.W.S., Chan, R.C.K., 2016. The neural transfer effect of working memory training to enhance hedonic processing in individuals with social anhedonia. Scientific Reports 6.

Lin, L., Chu, H., 2018. Quantifying publication bias in meta-analysis. Biometrics 74, 785–794.

Lipsey, M.W., 2003. Those confounded moderators in meta-analysis: Good, bad, and ugly. Ann Am Acad Polit Ss 587, 69–81.

Luo, Y., Eickhoff, S.B., Hetu, S., Feng, C.L., 2018. Social comparison in the brain: A coordinate-based meta-analysis of functional brain imaging studies on the downward and upward comparisons. Human Brain Mapping 39, 440–458.

Mahmoudi, A., Takerkart, S., Regragui, F., Boussaoud, D., Brovelli, A., 2012. Multivoxel Pattern Analysis for fMRI Data: A Review. Comput Math Method M.

Menon, V., Uddin, L.Q., 2010. Saliency, switching, attention and control: a network model of insula function. Brain Structure & Function 214, 655–667.

Monosov, I.E., 2017. Anterior cingulate is a source of valence-specific information about value and uncertainty. Nature Communications 8.

Muller, V.I., Cieslik, E.C., Laird, A.R., Fox, P.T., Radua, J., Mataix-Cols, D., Tench, C.R., Yarkoni, T., Nichols, T.E., Turkeltaub, P.E., Wager, T.D., Eickhoff, S.B., 2018. Ten simple rules for neuroimaging meta-analysis. Neuroscience and Biobehavioral Reviews 84, 151–161.

Naranjo, C.A., Tremblay, L.K., Busto, U.E., 2001. The role of the brain reward system in depression. Prog Neuropsychopharmacol Biol Psychiatry 25, 781–823.

Nasser, H.M., Calu, D.J., Schoenbaum, G., Sharpe, M.J., 2017. The Dopamine Prediction Error: Contributions to Associative Models of Reward Learning. Frontiers in Psychology 8.

Nawijn, L., van Zuiden, M., Koch, S.B., Frijling, J.L., Veltman, D.J., Olff, M., 2017. Intranasal oxytocin increases neural responses to social reward in post-traumatic stress disorder. Soc Cogn Affect Neurosci 12, 212–223.

Oldham, S., Murawski, C., Fornito, A., Youssef, G., Yucel, M., Lorenzetti, V., 2018a. The anticipation and outcome phases of reward and loss processing: A neuroimaging meta-analysis of the monetary incentive delay task. Hum Brain Mapp 39, 3398–3418.

Oldham, S., Murawski, C., Fornito, A., Youssef, G., Yucel, M., Lorenzetti, V., 2018b. The anticipation and outcome phases of reward and loss processing: A neuroimaging meta-analysis of the monetary incentive delay task. Human Brain Mapping 39, 3398–3418.

Ouzzani, M., Hammady, H., Fedorowicz, Z., Elmagarmid, A., 2016. Rayyan-a web and mobile app for systematic reviews. Systematic Reviews 5.

Piva, M., Velnoskey, K., Jia, R.N., Nair, A., Levy, I., Chang, S.W.C., 2019. The dorsomedial prefrontal cortex computes task-invariant relative subjective value for self and other. Elife 8.

Poldrack, R.A., Baker, C.I., Durnez, J., Gorgolewski, K.J., Matthews, P.M., Munafo, M.R., Nichols, T.E., Poline, J.B., Vul, E., Yarkoni, T., 2017. Scanning the horizon: towards transparent and reproducible neuroimaging research. Nature reviews. Neuroscience 18, 115–126.

Rademacher, L., Salama, A., Grunder, G., Spreckelmeyer, K.N., 2014. Differential patterns of nucleus accumbens activation during anticipation of monetary and social reward in young and older adults. Soc Cogn Affect Neurosci 9, 825–831.

Radua, J., Mataix-Cols, D., 2012. Meta-analytic methods for neuroimaging data explained. Biol Mood Anxiety Disord 2, 6.

Radua, J., Mataix-Cols, D., Phillips, M.L., El-Hage, W., Kronhaus, D.M., Cardoner, N., Surguladze, S., 2012. A new meta-analytic method for neuroimaging studies that combines reported peak coordinates and statistical parametric maps. European Psychiatry 27, 605–611.

Radua, J., Rubia, K., Canales-Rodriguez, E.J., Pomarol-Clotet, E., Fusar-Poli, P., Mataix-Cols, D., 2014. Anisotropic kernels for coordinate-based meta-analyses of neuroimaging studies. Front Psychiatry 5, 13.

Radua, J., Schmidt, A., Borgwardt, S., Heinz, A., Schlagenhauf, F., McGuire, P., Fusar-Poli, P., 2015. Ventral Striatal Activation During Reward Processing in Psychosis: A Neurofunctional Meta-Analysis. JAMA Psychiatry 72, 1243–1251.

Raichle, M.E., 2015. The Brain’s Default Mode Network. Annual Review of Neuroscience, Vol 38 38, 433–447.

Ramakrishnan, A., Byun, Y.W., Rand, K., Pedersen, C.E., Lebedev, M.A., Nicolelis, M.A.L., 2017. Cortical neurons multiplex reward-related signals along with sensory and motor information. Proceedings of the National Academy of Sciences of the United States of America 114, E4841–E4850.

Ramkumar, P., Dekleva, B., Cooler, S., Miller, L., Kording, K., 2016. Premotor and Motor Cortices Encode Reward. Plos One 11.

Ronchi, R., Bello-Ruiz, J., Lukowska, M., Herbelin, B., Cabrilo, I., Schaller, K., Blanke, O., 2015. Right insular damage decreases heartbeat awareness and alters cardio-visual effects on bodily self-consciousness. Neuropsychologia 70, 11–20.

Salomon, R., Levy, D.R., Malach, R., 2014. Deconstructing the Default: Cortical subdivision of the Default Mode/Intrinsic System During Self-Related Processing. Human Brain Mapping 35, 1491–1502.

Scheibner, H.J., Bogler, C., Gleich, T., Haynes, J.D., Bermpohl, F., 2017. Internal and external attention and the default mode network. Neuroimage 148, 381–389.

Schneider, M., Leuchs, L., Czisch, M., Samann, P.G., Spoormaker, V.I., 2018. Disentangling reward anticipation with simultaneous pupillometry/fMRI. Neuroimage 178, 11–22.

Shamseer, L., Moher, D., Clarke, M., Ghersi, D., Liberati, A., Petticrew, M., Shekelle, P., Stewart, L.A., Grp, P.-P., 2015. Preferred reporting items for systematic review and meta-analysis protocols (PRISMA-P) 2015: elaboration and explanation. Bmj-British Medical Journal 349.

Spreckelmeyer, K.N., Krach, S., Kohls, G., Rademacher, L., Irmak, A., Konrad, K., Kircher, T., Grunder, G., 2009. Anticipation of monetary and social reward differently activates mesolimbic brain structures in men and women. Soc Cogn Affect Neurosci 4, 158–165.

Spreckelmeyer, K.N., Rademacher, L., Paulus, F.M., Grunder, G., 2013a. Neural activation during anticipation of opposite-sex and same-sex faces in heterosexual men and women. Neuroimage 66, 223–231.

Spreckelmeyer, K.N., Rademacher, L., Paulus, F.M., Grunder, G., 2013b. Neural activation during anticipation of opposite-sex and same-sex faces in heterosexual men and women. Neuroimage 66, 223–231.

Spreng, R.N., 2012. The fallacy of a “task-negative” network. Frontiers in Psychology 3.

Sun, P., Zheng, L., Li, L., Guo, X.Y., Zhang, W.D., Zheng, Y.J., 2016. The Neural Responses to Social Cooperation in Gain and Loss Context. Plos One 11.

Sutton, B.P., Cheng, O.Y., Karampinos, D.C., Miller, G.A., 2009. Current trends and challenges in MRI acquisitions to investigate brain function. International Journal of Psychophysiology 73, 33–42.

Tanji, J., 1994. The supplementary motor area in the cerebral cortex. Neurosci Res 19, 251–268.

Todorov, A., 2012a. The role of the amygdala in face perception and evaluation. Motiv Emot 36, 16–26.

Todorov, A., 2012b. The role of the amygdala in face perception and evaluation. Motiv Emotion 36, 16–26.

Wang, D., Liu, T., Shi, J., 2017. Development of Monetary and Social Reward Processes. Sci Rep 7, 11128.

Wanke, N., Schwabe, L., 2019. Subjective Uncontrollability over Aversive Events Reduces Working Memory Performance and Related Large-Scale Network Interactions. Cereb Cortex.

Wilkos, E., Brown, T.J., Slawinska, K., Kucharska, K.A., 2015. Social cognitive and neurocognitive deficits in inpatients with unilateral thalamic lesions - pilot study. Neuropsychiatr Dis Treat 11, 1031–1038.

Wilson, R.P., Colizzi, M., Bossong, M.G., Allen, P., Kempton, M., Mtac, Bhattacharyya, S., 2018. The Neural Substrate of Reward Anticipation in Health: A Meta-Analysis of fMRI Findings in the Monetary Incentive Delay Task. Neuropsychol Rev 28, 496–506.

Wolff, M., Vann, S.D., 2019. The Cognitive Thalamus as a Gateway to Mental Representations. Journal of Neuroscience 39, 3–14.

Woo, C.W., Koban, L., Kross, E., Lindquist, M.A., Banich, M.T., Ruzic, L., Andrews-Hanna, J.R., Wager, T.D., 2014. Separate neural representations for physical pain and social rejection. Nat Commun 5, 5380.

Zheng, J., Anderson, K.L., Leal, S.L., Shestyuk, A., Gulsen, G., Mnatsakanyan, L., Vadera, S., Hsu, F.P.K., Yassa, M.A., Knight, R.T., Lin, J.J., 2017. Amygdala-hippocampal dynamics during salient information processing. Nature Communications 8.

Zhu, Y.J., Nachtrab, G., Keyes, P.C., Allen, W.E., Luo, L.Q., Chen, X.K., 2018. Dynamic salience processing in paraventricular thalamus gates associative learning. Science 362, 423–+.

