## Supplementary material for "Mapping social reward and punishment processing in the human brain: A voxel-based meta-analysis of neuroimaging findings using the Social Incentive Delay task"

**Table S1. Summary of sociodemographics and methodological characteristics of all included studies** (ASR – Anticipation of Social Reward; RSR – Receipt of Social Reward; ASP – Anticipation of Social Punishment Avoidance; RPR – Receipt of Social Punishment). In this table, we also detail which exact contrasts we could retrieve from each study (columns ASR, RSR, ASP and RSP) (please note that these columns refer to whether we could access the data needed for our meta-analysis and not whether the specific contrasts where used in a particular study; for some studies, some contrasts where used in the original study but we could not access the data we needed to include it in our meta-analysis)

| Author (year) | *N* | % Males | Mean Age | % Right-handed | Field strength | Spatial resolution | Smoothing | Software | Pharmacological study | ASR | RSR | ASP | RSP | Outcome type | Neutral feedback comparator | Variable incentive magnitude | Success rate |
|---|---|---|---|---|---|---|---|---|---|---|---|---|---|---|---|---|---|
| Barman et al. 2015 | 63 | 50.7 | 24.57 | NA | 3 | 2.5 x 2.5 x 3 | 8 | SPM8 | N/A | ✓ | ✓ | × | × | Visual (static faces) | White noise | × | Success rate set to 80% |
| Chan et al. 2016 | 28 | 28.6 | 18.88 | NA | 3 | 3.3 x 3.3 x 4 | 8 | SPM8 | N/A | ✓ | ✓ | ✓ | ✓ | Visual (static faces) | Neutral face | × | Reward: Success rate set to 66%<br><br>Punishment: Success rate set to 66% |
| Cremers et al. 2015 | 20 | 55 | 27.7 | NA | 3 | 2.75 x 2.75 x 2.75 | 6 | FSL 4.1.3 | N/A | ✓ | ✓ | ✓ | ✓ | Visual (static faces) | Morphed face | × | Reward: Success rate set to 66%<br><br>Punishment: Success rate set to 66% |
| Delmonte et al. 2012 | 21 | 100 | 17 | 100 | 3 | 3 x 3 x 3.5 | 5 | SPM8 | N/A | ✓ | ✓ | × | × | Visual (static faces) | Morphed face | × | Success rate set to 66% |
| Dutra et al. 2015 | 25 | 40 | 29.44 | 100 | 3 | 3.4 x 3.4 x 3.4 | 5 | FSL | N/A | ✓ | × | × | × | Praise (written) | Unspecified non-win outcome | ✓ | Success rate set to 50% |
| Goerlich et al. 2017 | 45 | 100 | 24.1 | 100 | 3 | 3.5 x 3.5 x 3.5 | 6 | SPM8 | N/A | ✓ | ✓ | × | × | Visual (dynamic) | Video depicting someone tapping fingers with eyes closed | × | Success rate set to 66% |
| Grope et al. 2013 | 14 | 0 | 26.1 | 100 | 3 | NA | 8 | SPM8 | Yes | ✓ | ✓ | × | × | Visual (static faces) | Dysmorphed faces | ✓ | Reward: Success rate set to 66%<br><br>Punishment: Success rate set to 66% |
| Kohls at al. 2013 | 22 | 50 | 25.6 | 100 | 3 | 3.5 x 3.5 x 3.5 | 5 | FSL 4.1.4 | N/A | ✓ | ✓ | ✓ | × | Visual (dynamic) | Video depicting neutral expression with natural body movement | × | Reward: Success rate set to 66%<br><br>Punishment: Success rate set to 66% |
| Nawinj et al. 2017 | 37 | 51 | 39.63 | NA | 3 | 3 x 3 x 3 | 8 | SPM8 | Yes | ✓ | ✓ | ✓ | ✓ | Visual (static faces) | Scrambled faces | × | Reward: Success rate set to 66%<br><br>Punishment: Success rate set to 34% |
| Richey et al. 2016 | 22 | 54.5 | 26.5 | 100 | 3 | 3.4 x 3.4 x 4 | 4 | FSL 4.1.4 | N/A | ✓ | ✓ | ✓ | ✓ | Visual (static faces) | Win or non-win symbol | × | Reward: Success rate set to 66% |

| | | | | | | | | | | | | | | | | | |
|---|---|---|---|---|---|---|---|---|---|---|---|---|---|---|---|---|---|
| | | | | | | | | | | | | | | | | | Punishment: Success rate set to 66% |
| Spreckelmeyer et al. 2013 | 30 | 56.7 | 22.9 | 100 | 3 | NA | 8 | SPM5 | N/A | ✓ | × | × | × | Visual (static faces) | Morphed face | ✓ | Success rate set to 66% |
| Utevsky et al. 2017 | 41 | 100 | 23.8 | NA | 3 | 3.8 x 3.8 x 3.8 | 5 | FSL 4.1.8 | N/A | ✓ | × | × | × | Visual (static faces) | Landscape | × | Success rate set to 60% |
| Gossen et al. 2014 | 35 | 100 | 24.085 | 100 | 3 | 3.5 x 3.5 x 3.5 | 6 | SPM8 | N/A | ✓ | × | × | × | Visual (dynamic) | Video depicting someone tapping fingers with eyes closed | × | Success rate set to 66% |
| Kirsch et al. 2003 | 27 | 11 | 23.3 | 100 | 1.5 | NA | 6 | SPM9 | N/A | ✓ | × | × | × | Verbal feedback (Fast/Slow) | Neutral symbol | × | Success rate around 63% |
| Stark et al. 2011 | 31 | 0 | 23 | 100 | 1.5 | NA | 9 | SPM8 | N/A | ✓ | × | × | × | Verbal feedback (Fast/Slow) | Neutral symbol | × | Success rate around 44.5% |
| Kollman et al. 2017 | 41 | 49 | 38.61 | NA | 3 | NA | 6.8 x 6.8 x 9 | SPM8 | N/A | ✓ | × | × | × | Verbal feedback (praise) | No feedback (did not require any response either) | × | NA |

**Table S2 – Anticipation of Social reward.** Meta-analytic results for the contrast anticipation of social reward versus anticipation of neutral feedback (16 studies, pooled sample size 502 participants). Results were considered significant for p<0.001, SDM-Z > 1 and cluster extent > 10 voxels as per current standard recommendations for multiple comparisons control using this method. The last column shows the result of the Egger's test for publication bias on each peak (p<0.05 suggests asymmetry of the funnel plot compatible with publication bias).

| Number | Description | MNI coordinates | Voxels | P | SDM-Z | Egger's test (p-value) |
|---|---|---|---|---|---|---|
| **Clusters ≥ 11 voxels with all voxels SDM-Z ≥ 2.705 and all peaks SDM-Z ≥ 2.948** | | | | | | |
| 1 | Left pons, right and left caudate and putamen, right thalamus, right olfactory cortex | 8,4,-6 | 2872 | 0 | 5.942 | 0.279 |
| 2 | Left precentral gyrus | -40,-16,56 | 1299 | $1.19x10^{-7}$ | 4.441 | 0.550 |
| 3 | Right supplementary motor area | 4,2,50 | 1217 | $1.19x10^{-7}$ | 4.435 | 0.019 |
| 4 | Left insula/frontal operculum | -32,14,10 | 370 | $7.15x10^{-7}$ | 4.267 | 0.044 |
| 5 | Paracingulate gyrus/ right superior frontal gyrus | 6,32,38 | 121 | $1.87x10^{-5}$ | 3.661 | 0.040 |
| 6 | Right precentral gyrus | 34,-8,46 | 92 | $3.11x10^{-5}$ | 3.549 | 0.055 |
| 7 | Precuneous | -16,-64,38 | 55 | $1.55x10^{-4}$ | 3.174 | 0.974 |
| 8 | Left lateral occipital gyrus, superior division | -24,-76,22 | 39 | $1.19x10^{-4}$ | 3.241 | 0.179 |
| 9 | Pons | 0,-32,-40 | 34 | $8.46x10^{-6}$ | 3.832 | 0.033 |
| 10 | Cerebellum, vermic lobule IV/V | 2,-50,-22 | 33 | $1.05x10^{-4}$ | 3.271 | 0.530 |
| 11 | Left precentral gyrus | -52,4,32 | 33 | $3.55x10^{-4}$ | 2.975 | 0.822 |
| 12 | Left lateral occipital gyrus, inferior division | -26,-88,6 | 24 | $1.90x10^{-4}$ | 3.129 | 0.027 |
| 13 | Right frontal orbital cortex/insular cortex | 28,26,-6 | 15 | $4.26x10^{-5}$ | 3.484 | 0.963 |
| 14 | Occipital pole/lateral occipital cortex | -26,-92,14 | 15 | $3.93x10^{-4}$ | 2.948 | 0.120 |
| 15 | Left inferior frontal gyrus, opercular part | -50,6,8 | 11 | $2.75x10^{-4}$ | 3.040 | 0.471 |
| **Clusters ≥ 10 voxels with all voxels SDM-Z ≤ -2.594 and all peaks SDM-Z ≤ -2.792** | | | | | | |
| 1 | Posterior cingulate | -4,-46,36 | 1519 | $2.98x10^{-7}$ | -3.837 | 0.156 |
| 2 | Right postcentral gyrus | 66,-10,28 | 1061 | 0 | -3.956 | 0.944 |
| 3 | Left middle temporal gyrus, posterior division | -56,-34,0 | 534 | $1.37x10^{-6}$ | -3.664 | 0.438 |
| 4 | Frontal pole | 48,48-4 | 468 | $5.36x10^{-7}$ | -3.790 | 0.554 |
| 5 | Left paracingulate gyrus/frontal medial cortex | -8,48,-6 | 410 | $1.132x10^{-6}$ | -3.720 | 0.594 |
| 6 | Right superior frontal gyrus, dorsolateral/frontal pole | 26,42,42 | 392 | $1.79x10^{-7}$ | -3.927 | 0.198 |
| 7 | Left middle frontal gyrus | -28,20,42 | 270 | $2.14x10^{-6}$ | -3.582 | 0.161 |
| 8 | Left cerebellum, crus I | -34,-70,-32 | 206 | $9.54x10^{-7}$ | -3.769 | 0.182 |
| 9 | Left angular gyrus | -46,-72,26 | 131 | $8.04x10^{-6}$ | -3.460 | 0.815 |
| 10 | Left middle temporal gyrus | -50,-10,-14 | 98 | $1.43x10^{-6}$ | -3.660 | 0.465 |
| 11 | Right middle temporal gyrus | 54,-66,20 | 63 | $9.65x10^{-5}$ | -3.063 | 0.586 |
| 12 | Left fusiform gyrus | -28,-30,-20 | 41 | $1.59x10^{-5}$ | -3.357 | 0.345 |
| 13 | Right Superior Frontal gyrus, medial | 12,60,36 | 31 | $1.57x10^{-4}$ | -2.972 | 0.339 |
| 14 | Left temporal fusiform gyrus | -28,-42,-16 | 28 | $8.36x10^{-5}$ | -3.088 | 0.873 |
| 15 | Right parahippocampal gyrus | 32,-22,-20 | 25 | $6.61x10^{-6}$ | -3.475 | 0.176 |
| 16 | Right middle temporal gyrus | 60,-62,0 | 20 | $1.91x10^{-5}$ | -3.327 | 0.993 |
| 17 | Left inferior frontal gyrus, triangular part | -54,20,26 | 20 | $5.51x10^{-5}$ | -3.162 | 0.342 |
| 18 | Right middle occipital gyrus | 46,-66,24 | 20 | $9.98x10^{-5}$ | -3.058 | 0.314 |
| 19 | Left inferior frontal gyrus, orbital part | -34,36,-10 | 18 | $5.57x10^{-5}$ | -3.160 | 0.873 |
| 20 | Right inferior frontal gyrus, orbital part | 30,38,-10 | 17 | $1.26x10^{-5}$ | -3.393 | 0.653 |
| 21 | Right superior frontal gyrus, dorsolateral | 22,60,14 | 17 | $1.95x10^{-4}$ | -2.933 | 0.905 |
| 22 | Right parahippocampal gyrus | 26,0,-30 | 11 | $1.60x10^{-4}$ | -2.968 | 0.574 |
| 23 | Left parietal operculum | -42,-18,22 | 10 | $5.66x10^{-5}$ | -3.156 | 0.245 |
| 24 | Right temporal fusiform cortex | 40,-40,-12 | 10 | $1.72x10^{-4}$ | -2.956 | 0.337 |
| 25 | Right superior frontal gyrus, medial | 8,48,4 | 10 | $3.96x10^{-4}$ | -2.792 | 0.632 |

**Table S3 – Receipt of Social Reward.** Meta-analytic results for the contrast receipt of social reward versus receipt of neutral feedback (9 studies, pooled sample size 272 participants). Results were considered significant for p<0.001, SDM-Z > 1 and cluster extent > 10 voxels as per current standard recommendations for multiple comparisons control using this method. The last column shows the result of the Egger's test for publication bias on each peak (p<0.05 suggests asymmetry of the funnel plot compatible with publication bias).

| Number | Description | MNI coordinates | Voxels | P | SDM-Z | Egger's test (p-value) |
|---|---|---|---|---|---|---|
| **Clusters ≥ 10 voxels with all voxels SDM-Z ≥ 3.059 and all peaks SDM-Z ≥ 3.319** | | | | | | |
| 1 | Right middle temporal gyrus | 52,-64,22 | 3592 | ~0 | 5.096 | 0.926 |
| 2 | Left middle occipital gyrus | -42,-68,2 | 2732 | $5.24x10^{-6}$ | 4.481 | 0.962 |
| 3 | Right temporal pole, superior temporal gyrus | 32,6,-20 | 2033 | ~0 | 6.241 | 0.190 |
| 4 | Right precentral gyrus | -36,6,-20 | 1194 | ~0 | 5.325 | 0.868 |
| 5 | Right calcarine fissure/surrounding cortex | -2,50,-18 | 790 | ~0 | 5.593 | 0.480 |
| 6 | Right superior frontal gyrus, medial | 6,60,16 | 601 | $2.11x10^{-5}$ | 4.161 | 0.734 |
| 7 | Left middle temporal gyrus | -46,-64,20 | 103 | $3.21x10^{-6}$ | 4.610 | 0.928 |
| 8 | Left middle temporal gyrus | -62,-50,8 | 62 | $4.51x10^{-5}$ | 3.954 | 0.125 |
| 9 | Left posterior cingulate gyrus | 0,-54,24 | 29 | $4.26x10^{-4}$ | 3.324 | 0.429 |
| 10 | Right lingual gyrus | 6,-32,-4 | 21 | $2.62x10^{-4}$ | 3.465 | 0.534 |
| 11 | Right inferior frontal gyrus, triangular part | 52,20,22 | 20 | $4.32x10^{-4}$ | 3.319 | 0.340 |
| 12 | Left middle temporal gyrus | -50,-50,12 | 15 | $8.83x10^{-5}$ | 3.777 | 0.809 |
| 13 | Left middle temporal gyrus | -58,-8,-14 | 14 | $2.65x10^{-4}$ | 3.464 | 0.719 |
| 14 | Left parahippocampal gyrus, posterior division | -22,-28,-18 | 11 | $2.14x10^{-4}$ | 3.551 | 0.984 |
| 15 | Right thalamus | 2,-10,-2 | 10 | $2.58x10^{-4}$ | 3.470 | 0.870 |
| 16 | Left frontal orbital cortex | -40,36,-8 | 10 | $1.46x10^{-4}$ | 3.635 | 0.545 |
| **Clusters ≥ 11 voxels with all voxels SDM-Z ≤ -3.275 and all peaks SDM-Z ≤ -3.543** | | | | | | |
| 1 | Precuneous | 20,-62,52 | 1626 | $2.14x10^{-6}$ | -4.248 | 0.203 |
| 2 | Right middle frontal gyrus | 26,4,50 | 504 | $4.17x10^{-7}$ | -4.332 | 0.647 |
| 3 | Right supramarginal gyrus | 62,-46,42 | 504 | $4.17x10^{-7}$ | -4.333 | 0.675 |
| 4 | Left superior parietal lobe/postcentral gyrus | -32,-40,40 | 212 | $4.76x10^{-6}$ | -4.193 | 0.422 |
| 5 | Right precuneus | 16,-56,18 | 178 | $1.19x10^{-7}$ | -4.438 | 0.376 |
| 6 | Left postcentral gyrus/supramarginal gyrus | -52,-22,28 | 180 | $1.25x10^{-6}$ | -4.282 | 0.133 |
| 7 | Left rolandic operculum, left heschl gyrus | -36,-34,20 | 179 | $1.59x10^{-5}$ | -4.069 | 0.136 |
| 8 | Left precuneus | -16,-58,14 | 159 | $1.19x10^{-7}$ | -4.455 | 0.415 |
| 9 | Left middle frontal gyrus | -22,0,50 | 151 | $2.98x10^{-7}$ | -4.344 | 0.280 |
| 10 | Right superior temporal gyrus | 56,-34,20 | 98 | $5.60x10^{-7}$ | -4.170 | 0.086 |
| 11 | Left rolandic operculum | -46,-10,8 | 67 | $3.94x10^{-5}$ | -3.932 | 0.111 |
| 12 | Right superior temporal gyrus | 60,-8,-4 | 53 | $1.96x10^{-6}$ | -4.259 | 0.423 |
| 13 | Left supramarginal gyrus | -54,-24,42 | 49 | $1.71x10^{-4}$ | -3.676 | 0.081 |
| 14 | Left supramarginal gyrus | -64,-32,36 | 27 | $1.63x10^{-5}$ | -4.064 | 0.120 |
| 15 | Right angular gyrus | 44,-74,42 | 23 | $1.91x10^{-4}$ | -3.654 | 0.488 |
| 16 | Right median cingulate/paracingulate gyri | 10,-6,44 | 20 | $9.94x10^{-5}$ | -3.780 | 0.056 |
| 17 | Right median cingulate/paracingulate gyri | 10,6,42 | 16 | $1.25x10^{-4}$ | -3.739 | 0.038 |
| 18 | Left middle occipital gyrus | -40,-80,38 | 14 | $7.23x10^{-5}$ | -3.839 | 0.126 |
| 19 | Left inferior parietal gyri | -62,-36,44 | 13 | $1.26x10^{-4}$ | -3.738 | 0.387 |
| 20 | Left precuneous | -8,-46,8 | 12 | $1.55x10^{-4}$ | -3.695 | 0.245 |
| 21 | Left insula | -46,-10,-6 | 12 | $3.36x10^{-4}$ | -3.543 | 0.049 |
| 22 | Right inferior parietal gyri | 52,-38,56 | 11 | $2.92x10^{-4}$ | -3.573 | 0.046 |

**Table S4 – Anticipation of Social Punishment Avoidance.** Meta-analytic results for the contrast anticipation of social punishment avoidance versus anticipation of neutral feedback (5 studies, pooled sample size 129 participants). Results were considered significant for p<0.001, SDM-Z > 1 and cluster extent > 10 voxels as per current standard recommendations for multiple comparisons control using this method. The last column shows the result of the Egger's test for publication bias on each peak (p<0.05 suggests asymmetry of the funnel plot compatible with publication bias).

| Number | Description | MNI coordinates | Voxels | P | SDM-Z | Egger's test (p-value) |
|---|---|---|---|---|---|---|
| **Clusters ≥ 10 voxels with all voxels SDM-Z ≥ 2.128 and all peaks SDM-Z ≥ 2.319** | | | | | | |
| 1 | Left occipital fusiform gyrus | -20,-88,-4 | 526 | $1.82x10^{-4}$ | 2.388 | 0.820 |
| 2 | Right striatum | 14,4,-4 | 448 | $5.96x10^{-7}$ | 3.452 | 0.501 |
| 3 | Left hippocampus, left thalamus | 2,-34,-6 | 151 | $1.78x10^{-4}$ | 2.390 | 0.836 |
| 4 | Right precentral gyrus | 52,6,38 | 140 | $1.80x10^{-4}$ | 2.390 | 0.806 |
| 5 | Right calcarine fissure/surrounding cortex | 16,-98,2 | 134 | $2.07x10^{-4}$ | 2.371 | 0.910 |
| 6 | Right supplementary motor area | 6,22,56 | 72 | $1.81x10^{-4}$ | 2.390 | 0.833 |
| 7 | Left precentral gyrus | -40,6,36 | 69 | $1.85x10^{-4}$ | 2.387 | 0.817 |
| 8 | Right frontal orbital cortex | 30,24,-16 | 59 | $1.70x10^{-4}$ | 2.400 | 0.957 |
| 9 | Left temporal, superior temporal gyrus | -42,22,-28 | 55 | $1.79x10^{-4}$ | 2.392 | 0.847 |
| 10 | Right middle temporal gyrus | 52,-4,-16 | 42 | $1.80x10^{-4}$ | 2.390 | 0.810 |
| 11 | Right middle temporal gyrus | 56,-42,10 | 36 | $1.82x10^{-4}$ | 2.389 | 0.877 |
| 12 | Right temporal occipital fusiform cortex | 32,-56,-10 | 36 | $1.82x10^{-4}$ | 2.388 | 0.875 |
| 13 | Left amygdala | -20,-10,-10 | 27 | $2.48x10^{-5}$ | 2.868 | 0.827 |
| 14 | Right middle temporal gyrus | 52,-70,0 | 20 | $2.14x10^{-4}$ | 2.368 | 0.926 |
| 15 | Posterior cingulate/ left parahippocampal gyrus, posterior division | 20,-36,0 | 19 | $1.92x10^{-4}$ | 2.380 | 0.905 |
| 16 | Left frontal orbital cortex | -40,36,-8 | 17 | $3.12x10^{-4}$ | 2.319 | 0.959 |
| 17 | Left middle temporal gyrus | -52,-20,-10 | 16 | $1.98x10^{-4}$ | 2.377 | 0.878 |
| 18 | Left striatum | -22,2,-2 | 15 | $5.10x10^{-5}$ | 2.700 | 0.426 |
| 19 | Right insula | 30,24,-4 | 10 | $1.30x10^{-4}$ | 2.465 | 0.972 |
| **Clusters ≥ 11 voxels with all voxels SDM-Z ≤ -2.323 and all peaks SDM-Z ≤ -2.366** | | | | | | |
| 1 | Left superior temporal gyrus | -58,-40,22 | 167 | $1.11x10^{-4}$ | -2.391 | 0.816 |
| 2 | Right postcentral gyrus | 54,-12,28 | 148 | $1.11x10^{-4}$ | -2.391 | 0.831 |
| 3 | Left inferior temporal gyrus | -54,-60,-10 | 129 | $1.29x10^{-4}$ | -2.389 | 0.814 |
| 4 | Left middle frontal gyrus | -34,46,14 | 97 | $9.87x10^{-5}$ | -2.393 | 0.843 |
| 5 | Right cerebellum VI | 26,-66,-24 | 86 | $1.16x10^{-4}$ | -2.391 | 0.824 |
| 6 | Left inferior parietal gyri | -26,-72,42 | 76 | $1.16x10^{-4}$ | -2.391 | 0.849 |
| 7 | Right superior frontal gyrus, medial orbital | 2,44,-2 | 52 | $1.16x10^{-4}$ | -2.390 | 0.856 |
| 8 | Left inferior parietal | -42,-42,48 | 38 | $1.22x10^{-4}$ | -2.390 | 0.857 |
| 9 | Right middle frontal gyrus | 34,38,42 | 31 | $1.43x10^{-4}$ | -2.387 | 0.822 |
| 10 | Left inferior frontal gyrus, opercular part | -58,6,18 | 31 | $1.91x10^{-4}$ | -2.380 | 0.783 |
| 11 | Left middle occipital gyrus | -26,-68,30 | 19 | $2.10x10^{-4}$ | -2.378 | 0.885 |
| 12 | Right frontal operculum | 42,4,12 | 17 | $1.04x10^{-4}$ | -2.392 | 0.853 |
| 13 | Left precuneus | -2,-60,50 | 14 | $1.43x10^{-4}$ | -2.387 | 0.820 |
| 14 | Left fusiform gyrus | -26,-42,-12 | 12 | $1.60x10^{-4}$ | -2.385 | 0.834 |
| 15 | Right middle frontal gyrus | 46,38,30 | 12 | $1.99x10^{-4}$ | -2.380 | 0.866 |
| 16 | Right angular gyrus | 42,-70,34 | 12 | $2.85x10^{-4}$ | -2.371 | 0.770 |
| 17 | Left cuneus cortex | -16,-74,32 | 12 | $1.60x10^{-4}$ | -2.384 | 0.874 |
| 18 | Left superior temporal gyrus/planum temporale | -52,-34,6 | 12 | $2.32x10^{-4}$ | -2.376 | 0.911 |
| 19 | Right middle frontal gyrus | 24,46,30 | 12 | $3.42x10^{-4}$ | -2.366 | 0.823 |
| 20 | Left temporal pole, superior temporal gyrus, BA 38 | -58,6,-2 | 11 | $2.97x10^{-4}$ | -2.370 | 0.748 |

**Table S5 – Receipt of Social Punishment.** Meta-analytic results for the contrast receipt of social punishment versus receipt of neutral feedback (4 studies, pooled sample size 107 participants). Results were considered significant for p<0.001, SDM-Z > 1 and cluster extent > 10 voxels as per current standard recommendations for multiple comparisons control using this method. The last column shows the result of the Egger's test for publication bias on each peak ($p<0.05$ suggests asymmetry of the funnel plot compatible with publication bias).

| Number | Description | MNI coordinates | Voxels | P | SDM-Z | Egger's test (p-value) |
|---|---|---|---|---|---|---|
| **Clusters ≥ 10 voxels with all voxels SDM-Z ≥ 2.961 and all peaks SDM-Z ≥ 3.439** | | | | | | |
| 1 | Right inferior temporal gyrus, BA 19 | 50,-72,-8 | 582 | $2.05 \times 10^{-5}$ | 4.446 | 0.867 |
| 2 | Right inferior frontal gyrus, triangular part | 50,34,4 | 385 | $6.19 \times 10^{-6}$ | 4.916 | 0.898 |
| 3 | Left inferior frontal gyrus, orbital part | -40,28,-4 | 126 | $2.09 \times 10^{-5}$ | 4.436 | 0.842 |
| 4 | Left inferior frontal gyrus, triangular part | -48,22,22 | 74 | $3.54 \times 10^{-5}$ | 4.243 | 0.592 |
| 5 | Left inferior frontal gyrus/frontal operculum | -36,24,12 | 64 | $7.50 \times 10^{-5}$ | 3.968 | 0.863 |
| 6 | Right temporal pole, middle temporal gyrus | 58,8,-20 | 22 | $2.55 \times 10^{-5}$ | 4.368 | 0.866 |
| 7 | Right superior frontal gyrus, medial | 6,52,30 | 19 | $4.66 \times 10^{-5}$ | 4.151 | 0.851 |
| 8 | Left temporal pole | -40,12,-32 | 14 | $7.53 \times 10^{-5}$ | 3.965 | 0.862 |
| 9 | Right superior frontal gyrus, medial | 6,52,46 | 15 | $2.80 \times 10^{-4}$ | 3.439 | 0.997 |
| 10 | Left frontal orbital cortex | -46,26,-14 | 10 | $1.90 \times 10^{-4}$ | 3.598 | 0.749 |
| 11 | Left frontal orbital cortex/frontal pole | -34,34,-18 | 10 | $2.46 \times 10^{-4}$ | 3.496 | 0.736 |
| **Clusters ≥ 10 voxels with all voxels SDM-Z ≤ -2.460 and all peaks SDM-Z ≤ -2.647** | | | | | | |
| 1 | Left cuneus cortex | 0,-82,32 | 190 | $5.84 \times 10^{-5}$ | -2.735 | 0.609 |
| 2 | Left angular gyrus | -46,-64,44 | 185 | $5.41 \times 10^{-5}$ | -2.737 | 0.616 |
| 3 | Right angular gyrus | 54,-54,40 | 74 | $4.69 \times 10^{-5}$ | -2.742 | 0.629 |
| 4 | Right supramarginal gyrus | 52,-40,52 | 38 | $5.84 \times 10^{-5}$ | -2.735 | 0.618 |
| 5 | Left superior/middle frontal gyrus | -26,18,56 | 37 | $4.69 \times 10^{-5}$ | -2.742 | 0.654 |
| 6 | Right superior frontal gyrus, dorsolateral | 30,2,60 | 38 | $1.30 \times 10^{-4}$ | -2.699 | 0.725 |
| 7 | Frontal pole | -22,66,14 | 37 | $5.84 \times 10^{-5}$ | -2.736 | 0.648 |
| 8 | Left superior frontal gyrus/frontal pole | -20,38,42 | 31 | $2.56 \times 10^{-4}$ | -2.647 | 0.790 |
| 9 | Right superior parietal gyrus | 20,-68,62 | 30 | $4.83 \times 10^{-5}$ | -2.740 | 0.639 |
| 10 | Left caudate nucleus | -10,16,4 | 26 | $7.29 \times 10^{-5}$ | -2.728 | 0.641 |
| 11 | Right superior frontal gyrus, dorsolateral | 34,60,10 | 24 | $5.84 \times 10^{-5}$ | -2.736 | 0.687 |
| 12 | Left superior frontal gyrus, dorsolateral | -20,32,52 | 21 | $5.23 \times 10^{-5}$ | -2.738 | 0.656 |
| 13 | Right insula | 32,-12,14 | 18 | $6.10 \times 10^{-5}$ | -2.735 | 0.632 |
| 14 | Right middle temporal gyrus | 54,-34,-14 | 17 | $1.79 \times 10^{-4}$ | -2.677 | 0.762 |
| 15 | Right superior temporal gyrus | 68,-16,8 | 13 | $7.14 \times 10^{-5}$ | -2.728 | 0.673 |
| 16 | Left frontal pole | -28,52,2 | 12 | $4.83 \times 10^{-5}$ | -2.740 | 0.666 |
| 17 | Left middle frontal gyrus | -44,14,46 | 11 | $1.10 \times 10^{-4}$ | -2.709 | 0.567 |
| 18 | Left middle/inferior frontal gyrus | -44,26,38 | 10 | $1.23 \times 10^{-4}$ | -2.701 | 0.703 |

**Fig. S1 – Anticipation of Social reward (Jackknife sensitivity analysis).** Colour bars represent the number of binarized jackknife maps including increases (red) or decreases (blue) in BOLD for this contrast in a certain voxel. For reference, the reader can think of these number as the higher the number of maps where a certain voxel is included, the higher the robustness of its involvement in this contrast.

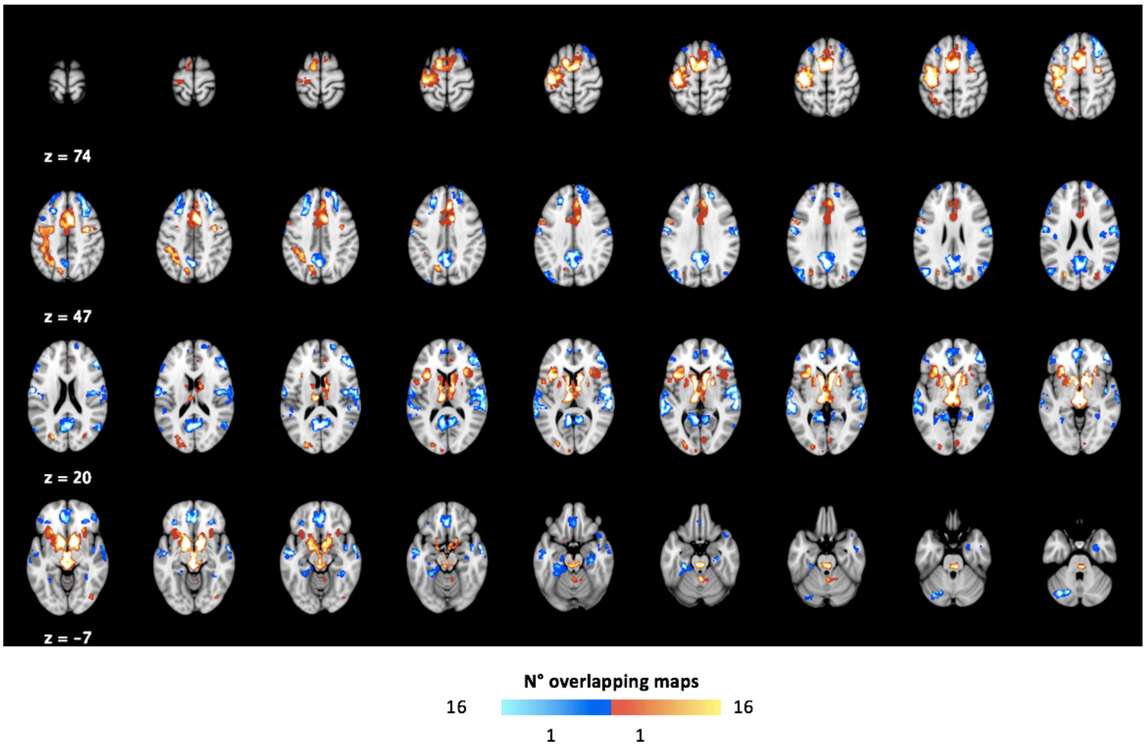

**Fig. S2 – Anticipation of Social reward (Heterogeneity).** In this figure, we present areas where we found considerable heterogeneity across studies as assessed by the $I^2$ statistics. To identify areas of relevant heterogeneity, we thresholded these maps for $I^2$>40% and masked them to retain only voxels where we found significant increases/decreases in BOLD in our main meta-analyses for this contrast.

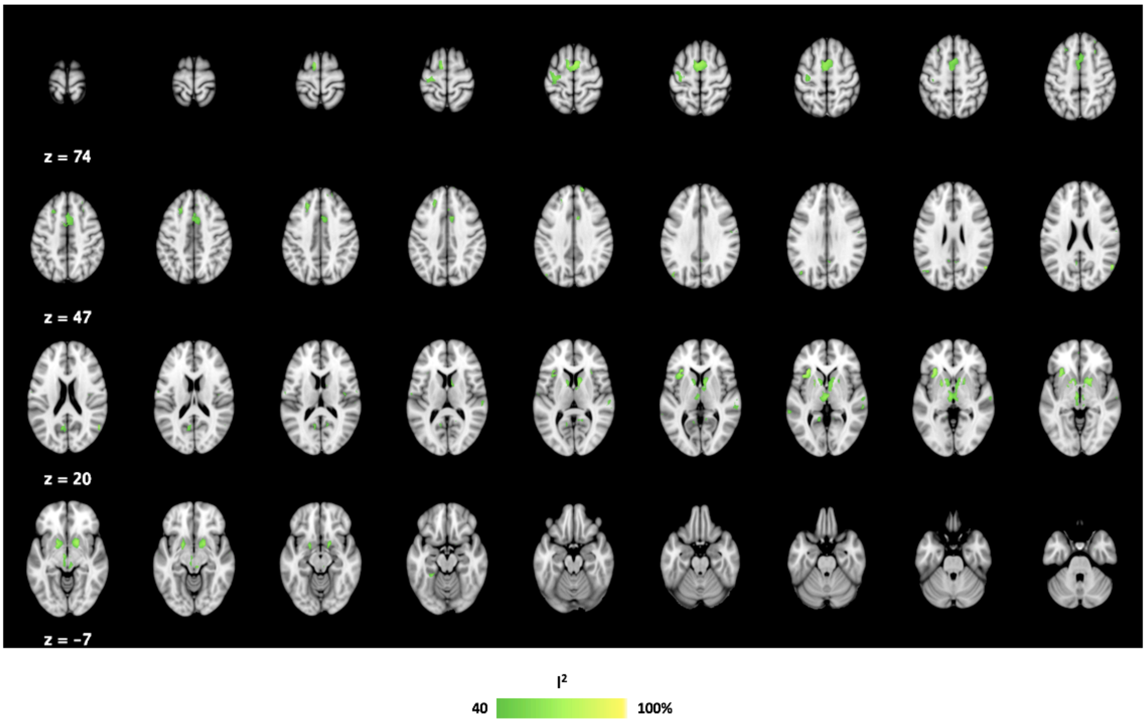


**Fig. S3 – Anticipation of Social reward (subgroup analysis – without studies using verbal feedback).** In this figure, we present the results of a subgroup analysis conducted after excluding all studies using verbal feedback as outcome. Results were considered significant for p<0.001, Z-SDM > 1 and cluster extent > 10 voxels as per current standard recommendations for multiple comparisons control using this method.

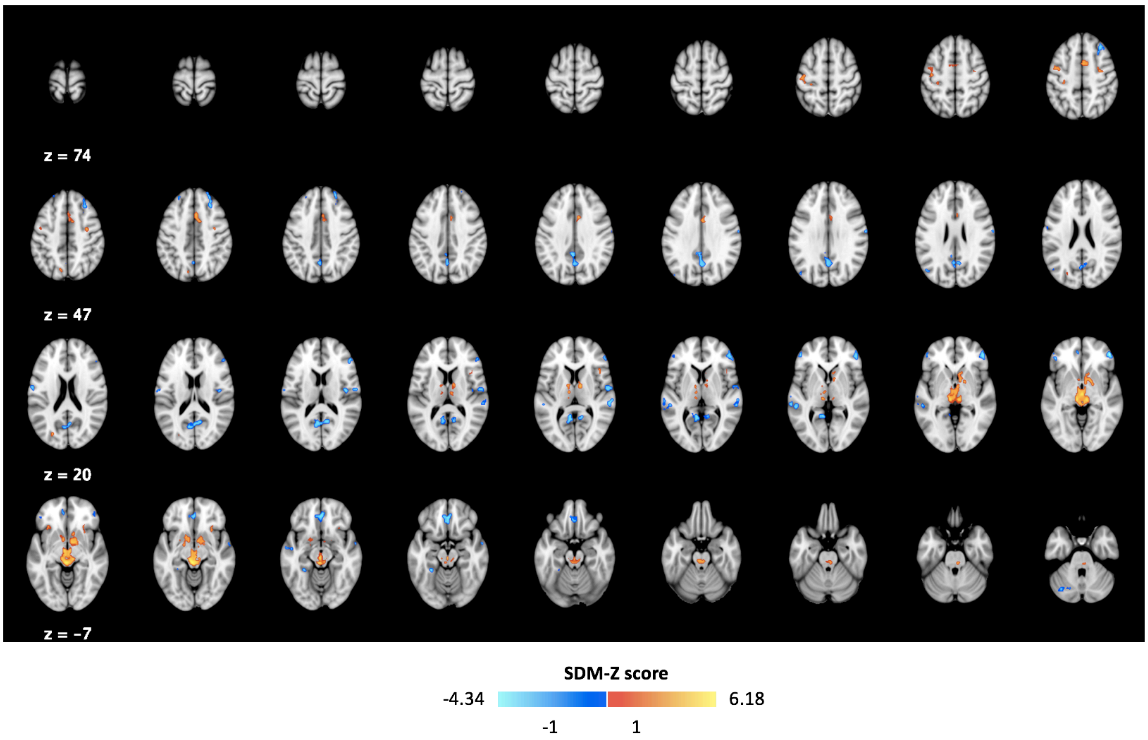


**Fig. S4 – Anticipation of Social reward (subgroup analysis – only studies using static emotional faces).** In this figure, we present the results of a subgroup analysis conducted using only studies where static emotional faces were presented as outcome. Results were considered significant for p<0.001, SDM-Z > 1 and cluster extent > 10 voxels as per current standard recommendations for multiple comparisons control using this method.

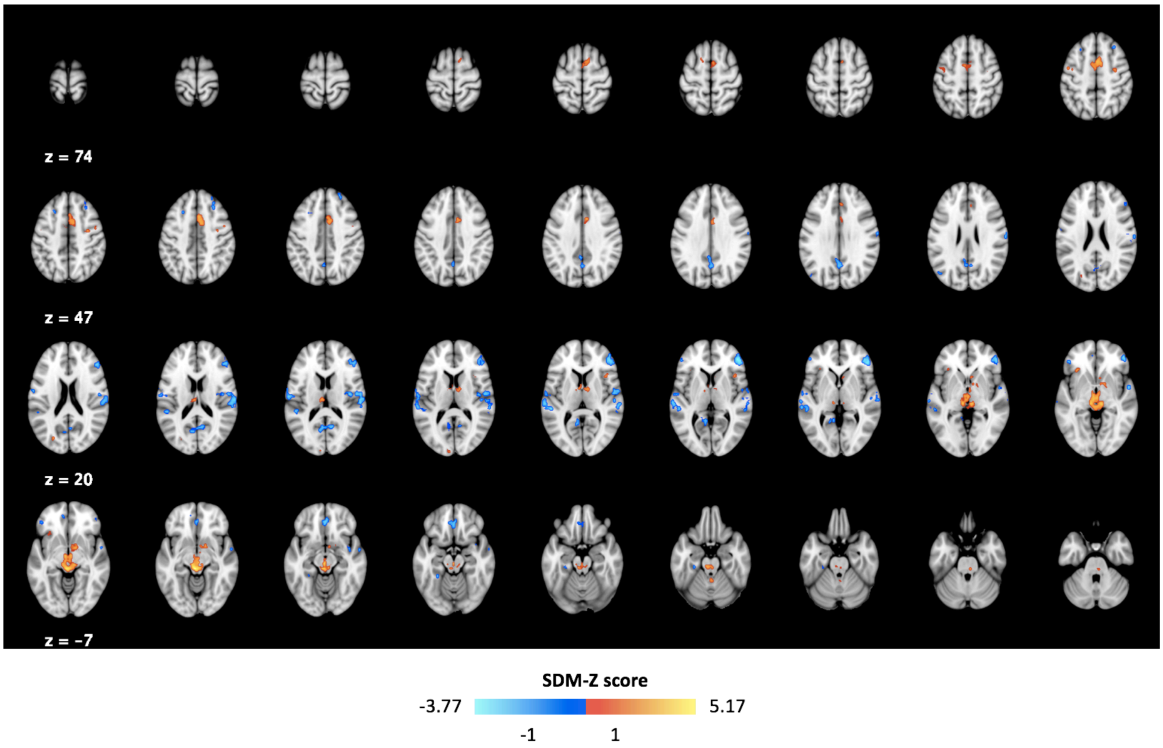


**Fig. S5 – Anticipation of Social reward (subgroup analysis – without placebo).** In this figure, we present the results of a subgroup analysis conducted after excluding two studies where administration of an intranasal placebo was performed. Results were considered significant for p<0.001, SDM-Z > 1 and cluster extent > 10 voxels as per current standard recommendations for multiple comparisons control using this method.

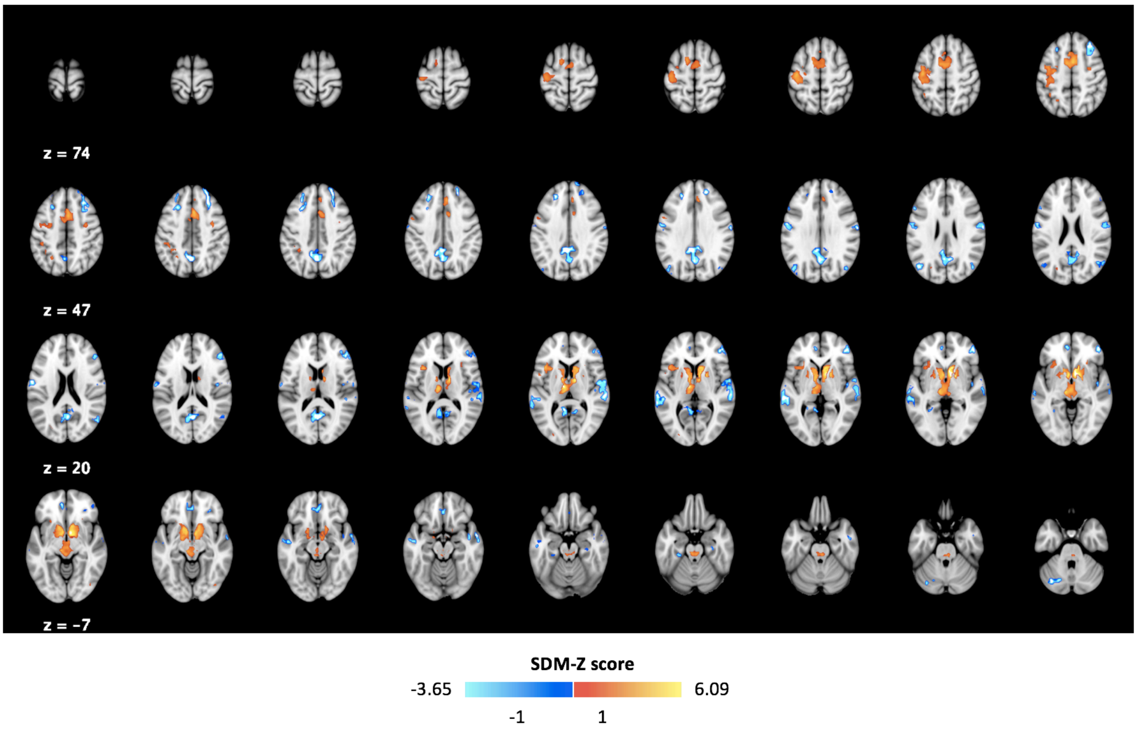

**Fig. S6 – Receipt of Social reward (Jackknife sensitivity analysis).** Colour bars represent the number of binarized jackknife maps including increases (red) or decreases (blue) in BOLD for this contrast in a certain voxel.

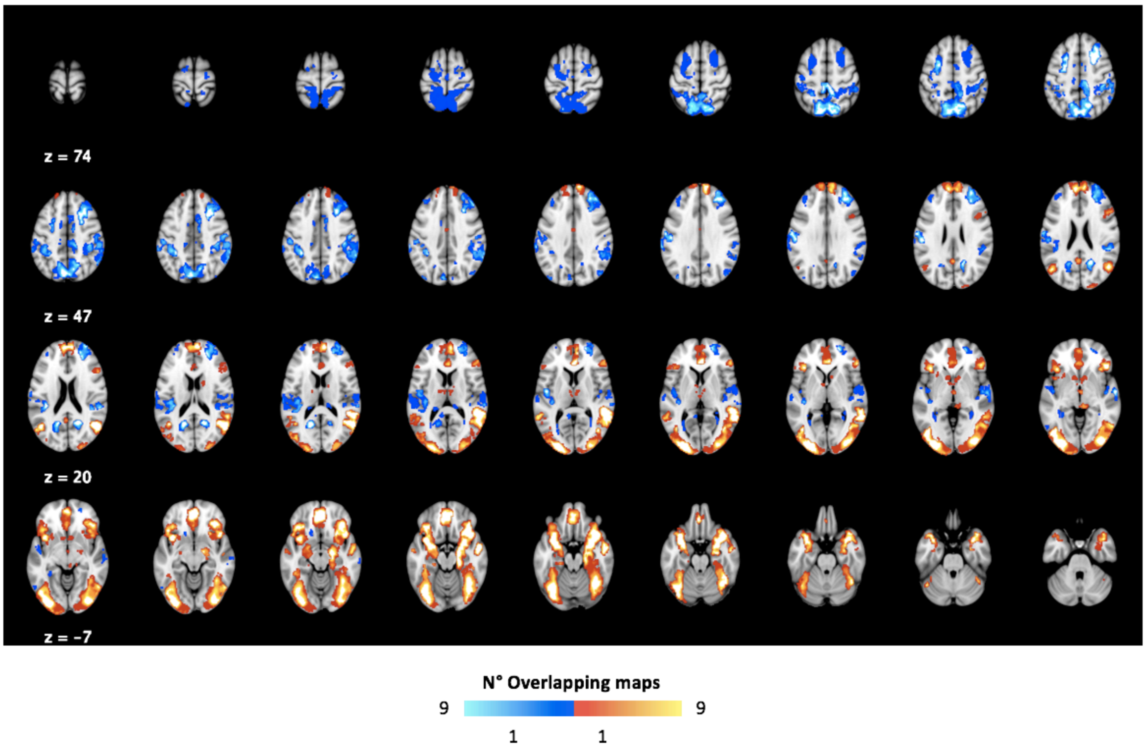


**Fig. S7 – Receipt of Social reward feedback (Heterogeneity).** In this figure, we present areas where we found considerable heterogeneity across studies as assessed by the $I^2$ statistics. To identify areas of relevant heterogeneity, we thresholded these maps for $I^2$>40% and masked them to retain only voxels where we found significant increases/decreases in BOLD in our main meta-analyses for this contrast.

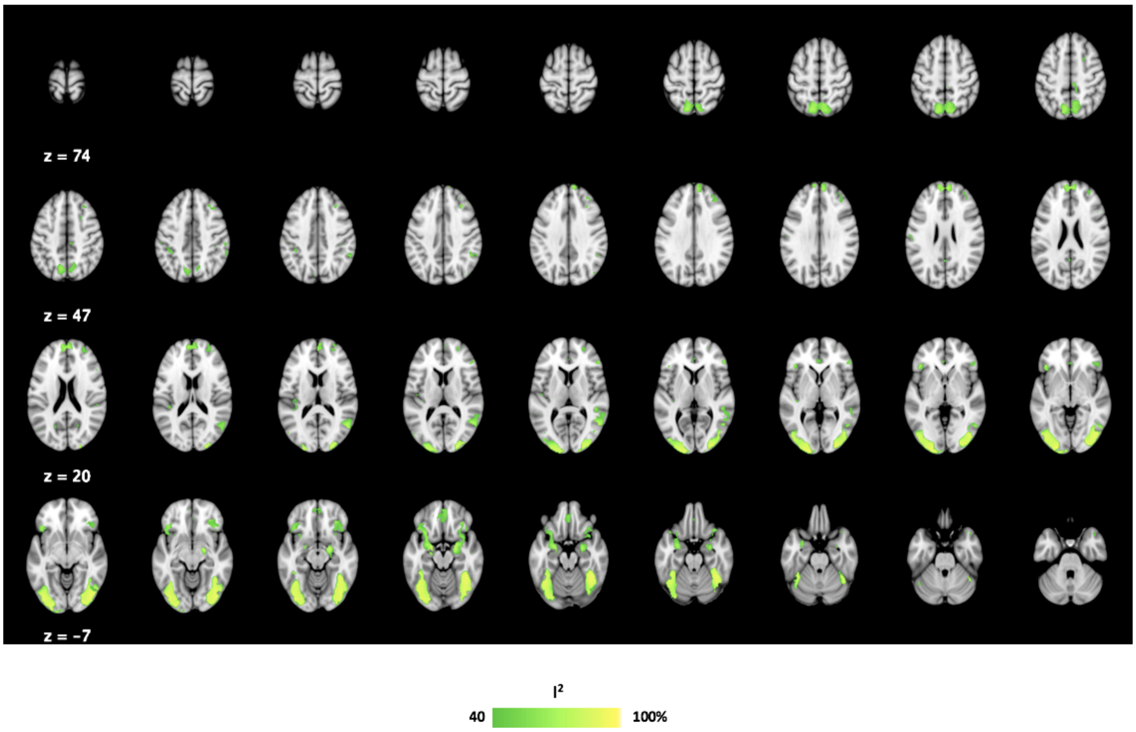


**Fig. S8 – Receipt of Social reward (subgroup analysis – only studies using static emotional faces).** In this figure, we present the results of a subgroup analysis conducted using only studies where static emotional faces were presented as outcome. Results were considered significant for $p<0.001$, SDM-Z > 1 and cluster extent > 10 voxels as per current standard recommendations for multiple comparisons control using this method.

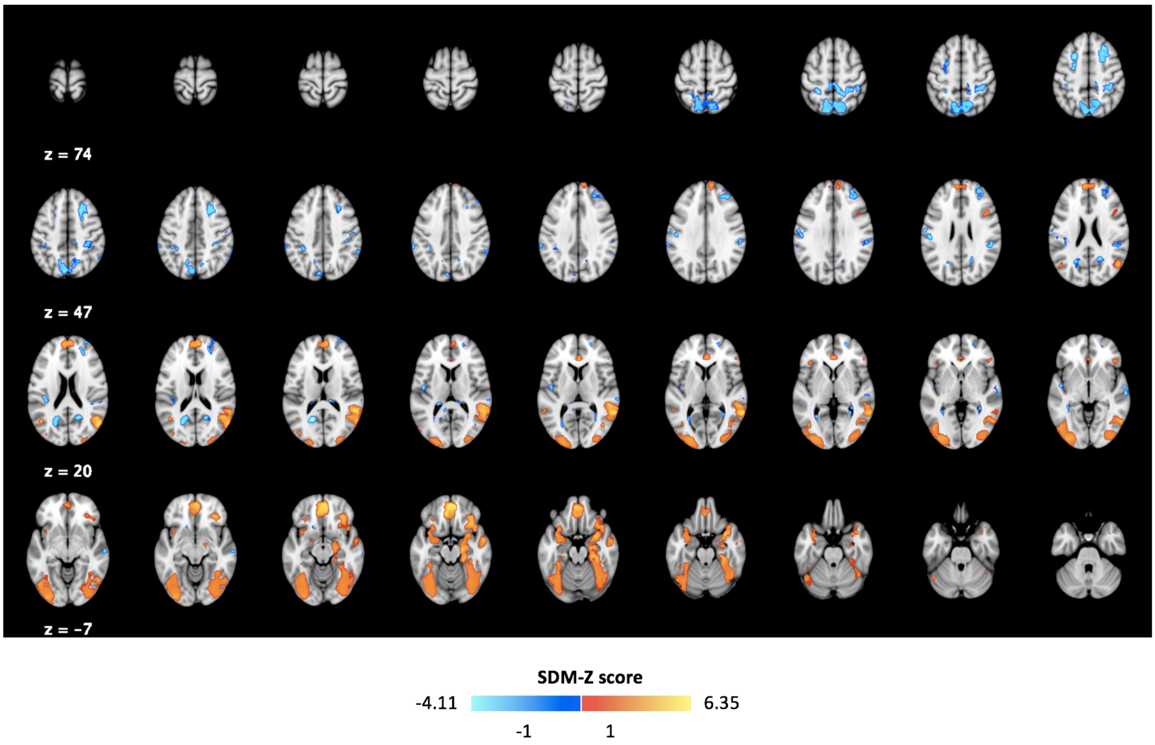


**Fig. S9 – Receipt of Social reward (subgroup analysis – without pharmacological studies).** In this figure, we present the results of a subgroup analysis conducted after excluding two studies where administration of an intranasal placebo was performed. Results were considered significant for $p<0.001$, SDM-Z > 1 and cluster extent > 10 voxels as per current standard recommendations for multiple comparisons control using this method.

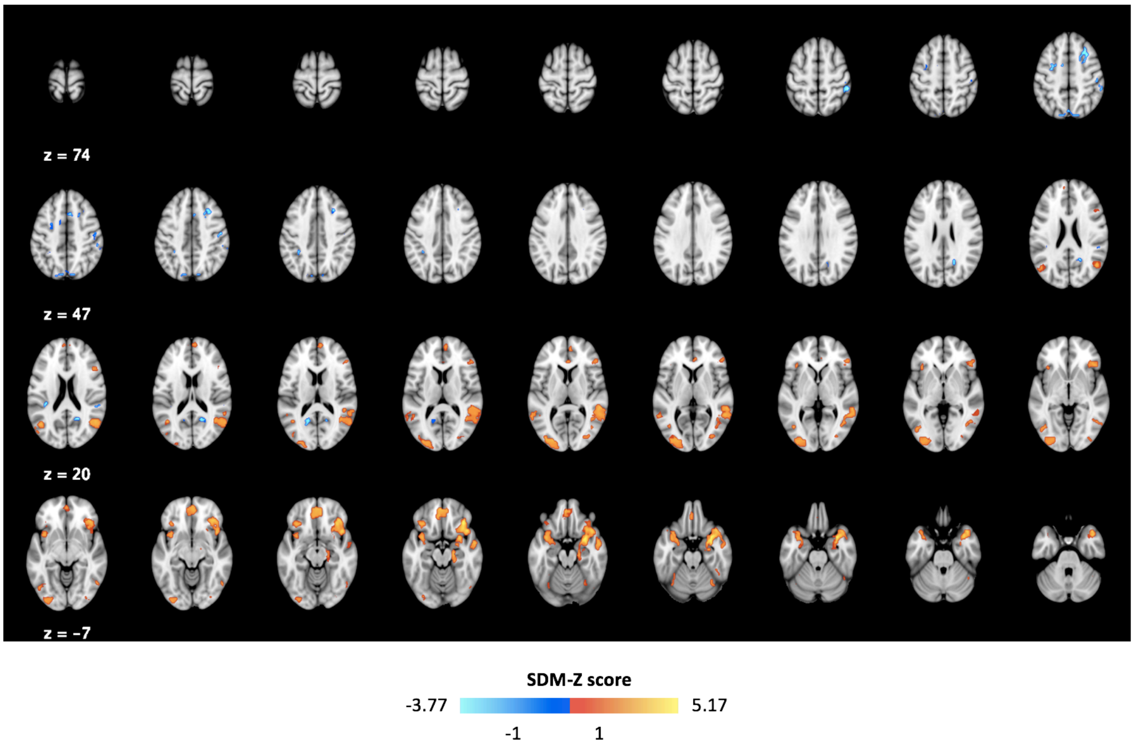

**Fig. S10 – Anticipation of Social punishment avoidance (Jackknife sensitivity analysis).** Colour bars represent the number of binarized jackknife maps including increases (red) or decreases (blue) in BOLD for this contrast in a certain voxel.

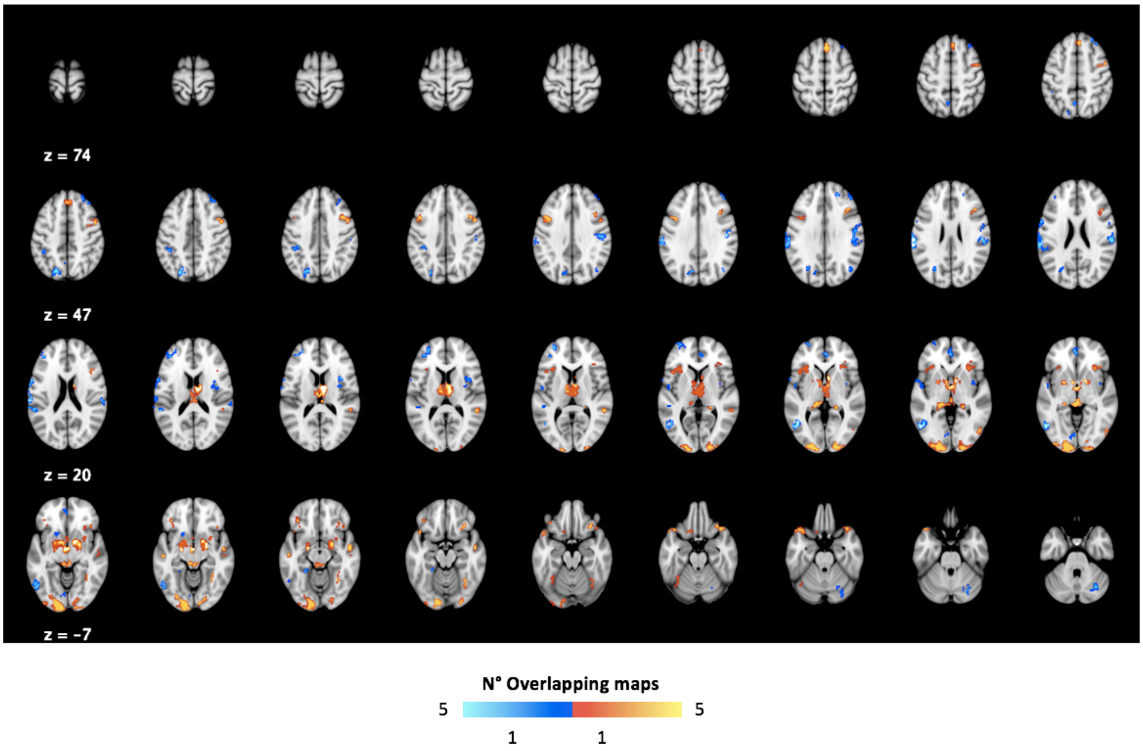


**Fig. S11 – Anticipation of Social punishment avoidance (Heterogeneity).** In this figure, we present areas where we found considerable heterogeneity across studies as assessed by the $I^2$ statistics. To identify areas of relevant heterogeneity, we thresholded these maps for $I^2>40\%$ and masked them to retain only voxels where we found significant increases/decreases in BOLD in our main meta-analyses for this contrast.

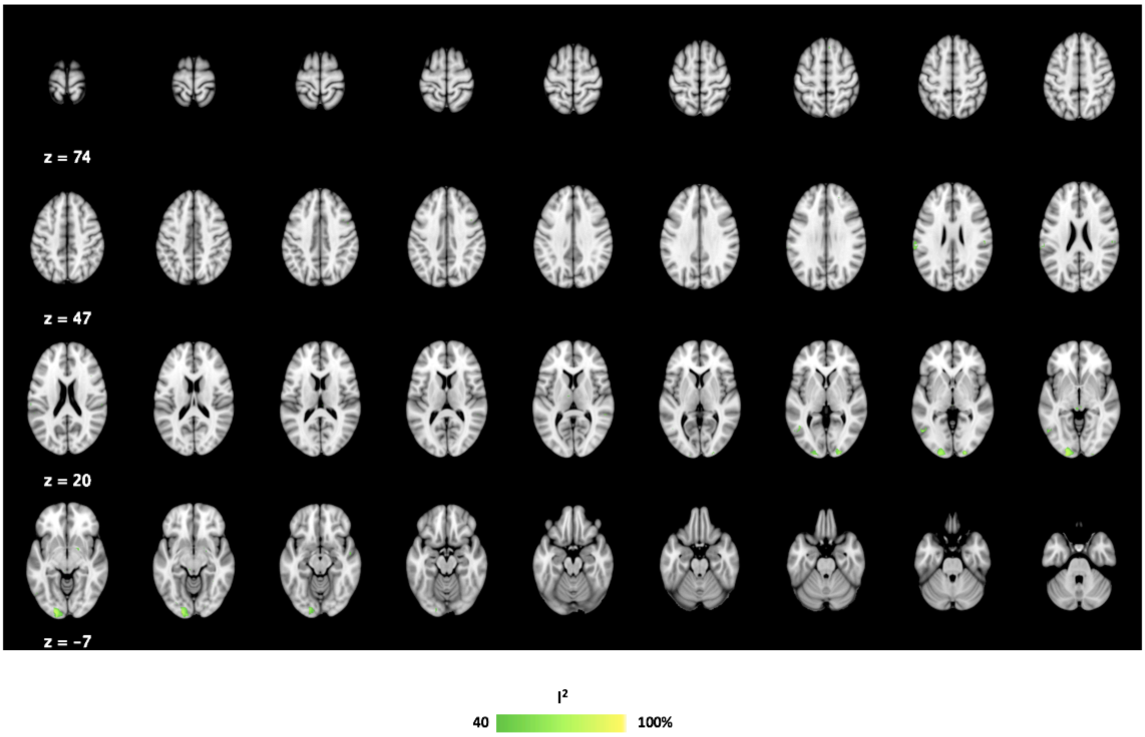


**Fig. S12 – Anticipation of Social punishment avoidance (Metaregression - gender).** In this figure, we present the results of a metaregression analysis for the percentage of male participants included in each study. Results were considered significant for $p<0.0005$ (0.001/2, since we conducted 2 metaregressions), SDM-Z > 1 and cluster extent > 10 voxels as per current standard recommendations for multiple comparisons control using this method.

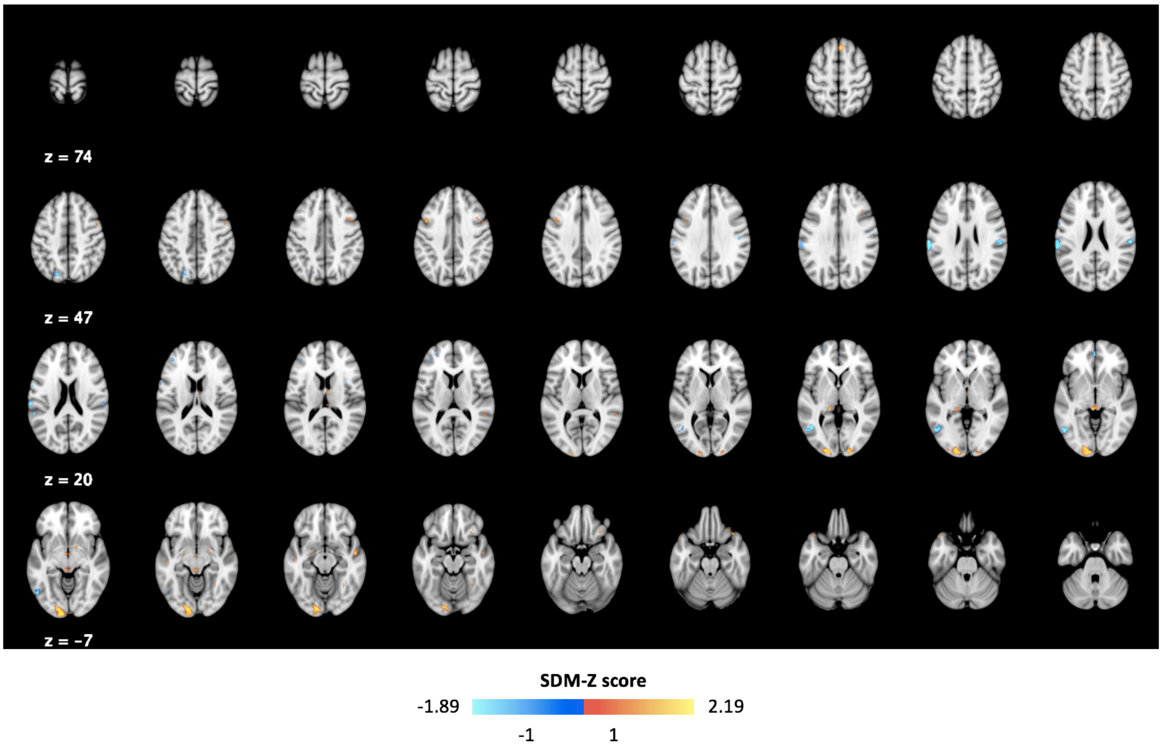


**Fig. S13 – Receipt of Social punishment (Jackknife sensitivity analysis).** Colour bars represent the number of binarized jackknife maps including increases (red) or decreases (blue) in BOLD for this contrast in a certain voxel.

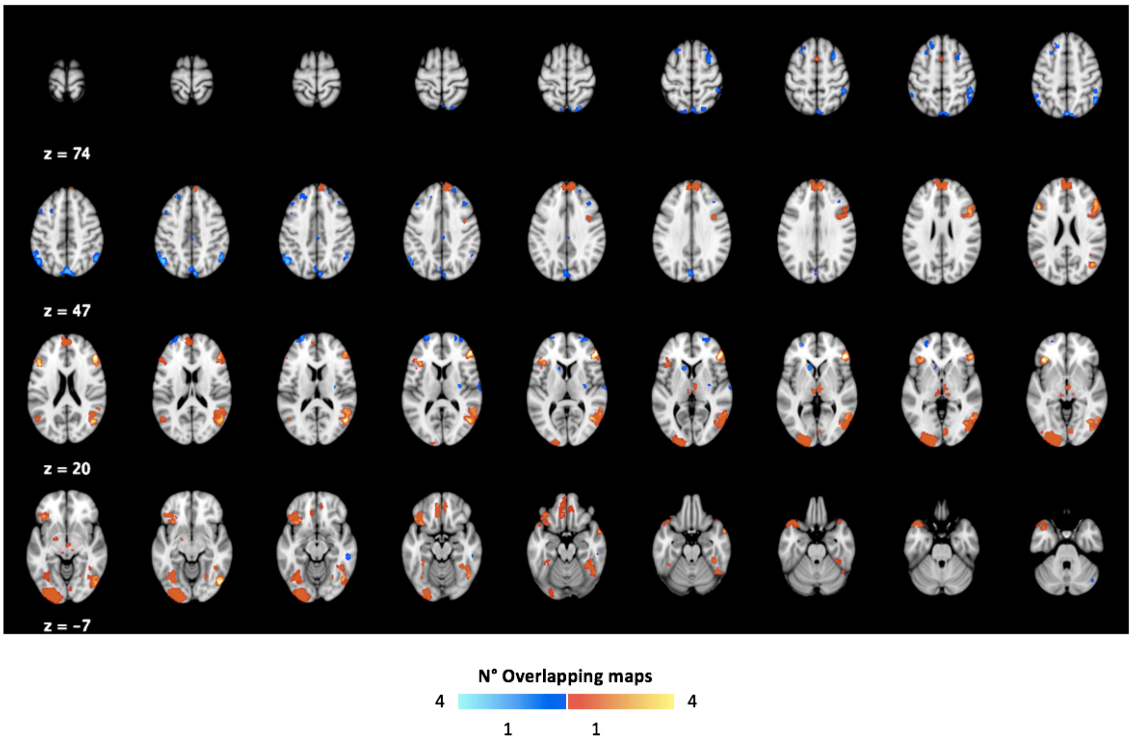

**Fig. S14 – Receipt of Social punishment (Heterogeneity).** In this figure, we present areas where we found considerable heterogeneity across studies as assessed by the $I^2$ statistics. To identify areas of relevant heterogeneity, we thresholded these maps for $I^2$>40% and masked them to retain only voxels where we found significant increases/decreases in BOLD in our main meta-analyses for this contrast.

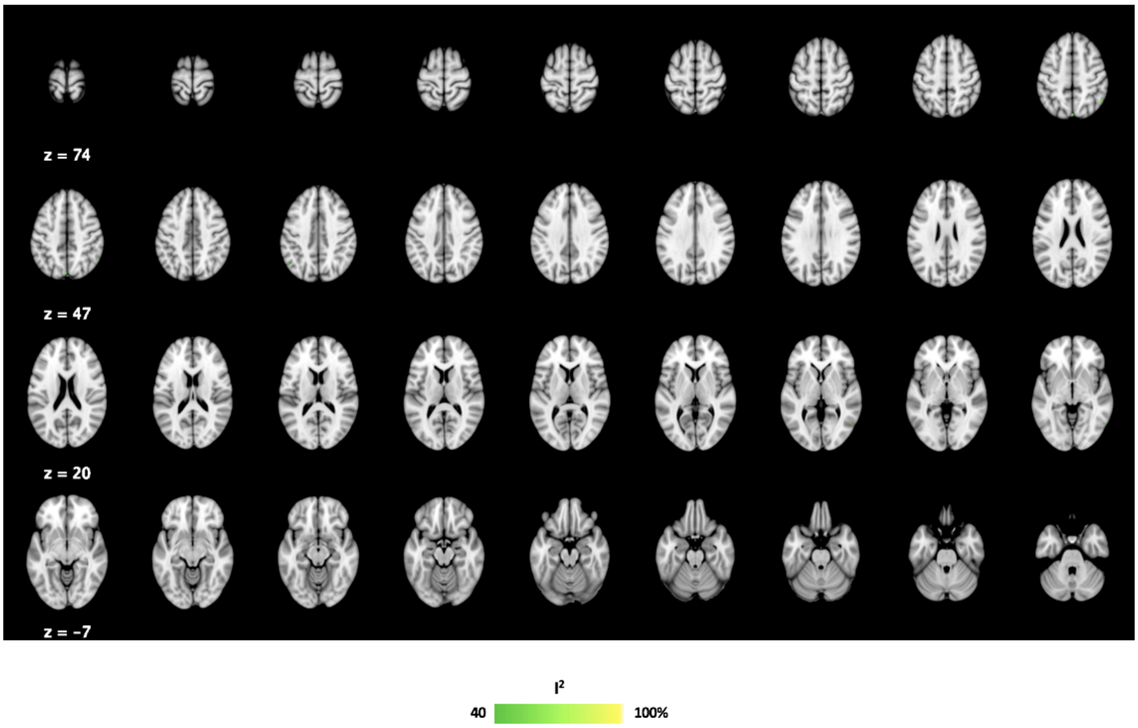


**Fig. S15 – Overview of the brain coverage of the data included in each of our four meta-analysis.** A. Anticipation of Social Reward; B. Receipt of Social Reward; C. Anticipation of Social Punishment; D. Receipt of Social Punishment. The colour bar represents the number of studies containing data for each voxel.

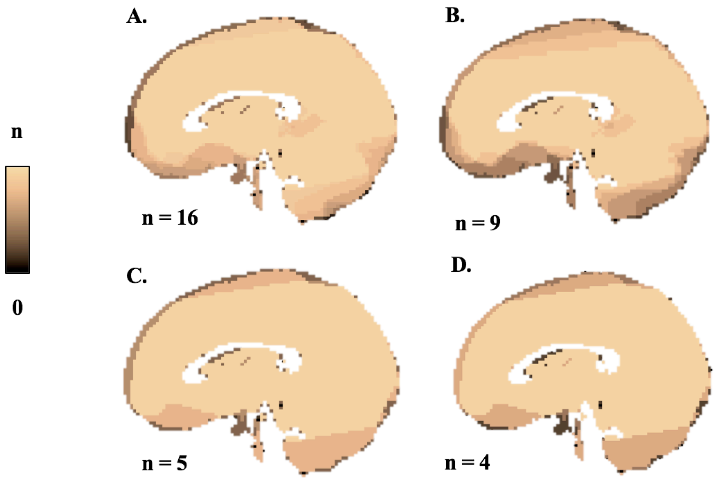

**Fig. S16 – Adjusted analysis accounting for dropout in the ventromedial prefrontal cortex**. A. Anticipation of Social Reward; B. Receipt of Social Reward; C. Anticipation of Social Punishment avoidance; D. Receipt of Social Punishment. In this figure, we thresholded both the original and adjusted maps using the average of the critical SDM-Z values generated in the permutation tests (50 permutations) for the original analyses.

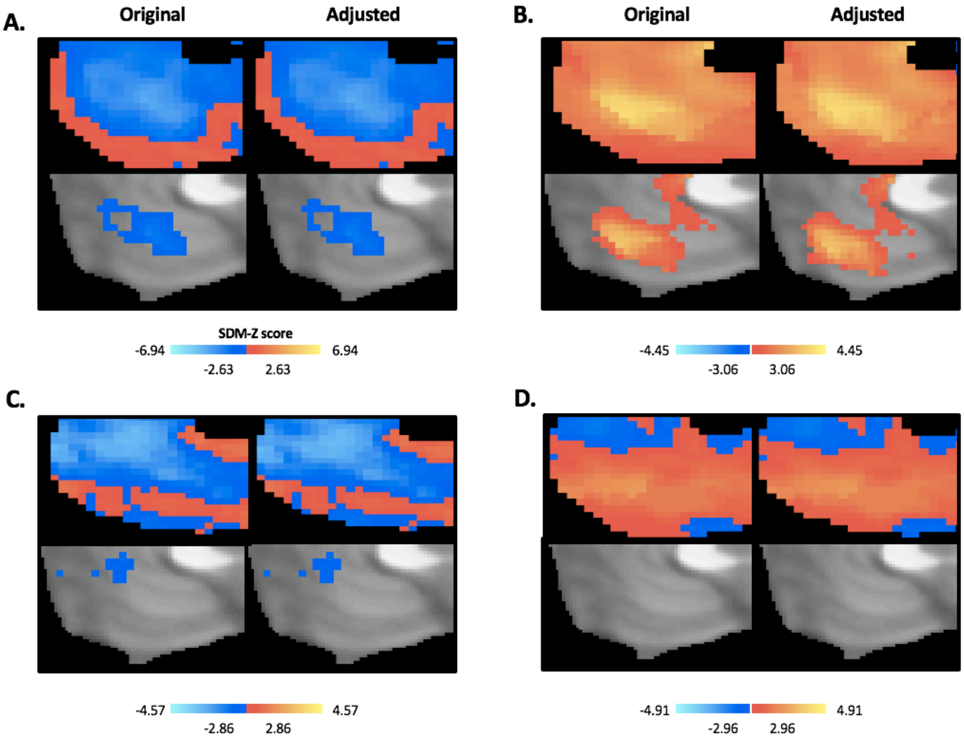


**Fig. S17 – Anticipation of Social AND monetary rewards.** In this figure, we provide a co-representation of our meta-analytic map for the anticipation of social reward (green) and a meta-analytic map for the contrast anticipation of monetary reward versus anticipation of neutral feedback (red) retrieved from a previous meta-analysis. We thresholded both maps using the same criteria: $p<0.001$, SDM-Z > 1 and cluster extent > 10 voxels and then binarize them. Voxels belonging to both maps are coloured in yellow.

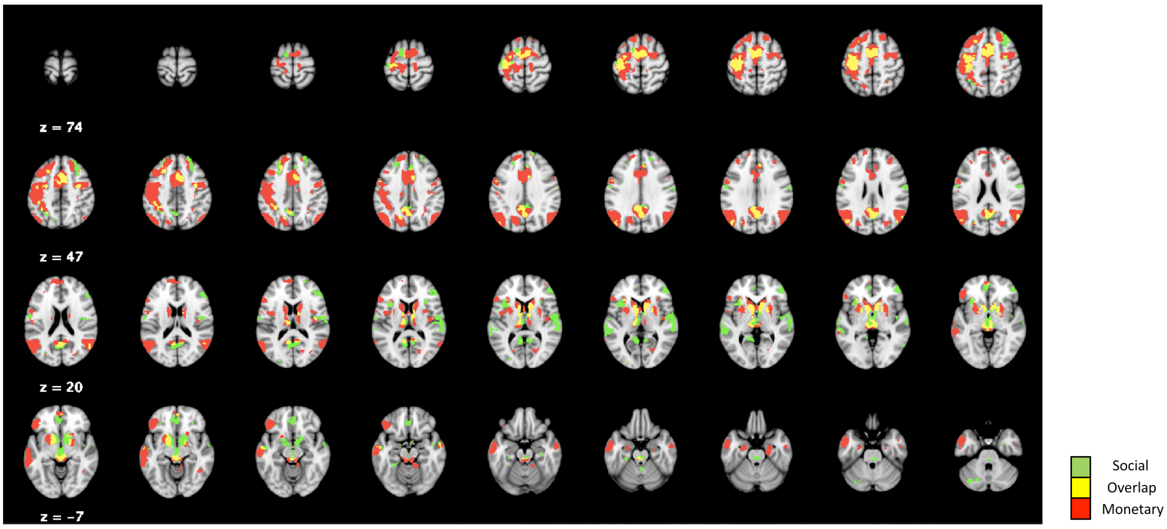


**Fig. S18 – Anticipation of Social AND monetary punishments avoidance.** In this figure, we provide a co-representation of our meta-analytic map for the anticipation of social punishment avoidance (green) and a meta-analytic map for the contrast anticipation of monetary loss avoidance versus anticipation of neutral feedback (red) retrieved from a previous meta-analysis. We thresholded both maps using the same criteria: p<0.001, SDM-Z > 1 and cluster extent > 10 voxels and then binarize them. Voxels belonging to both maps are coloured in yellow.

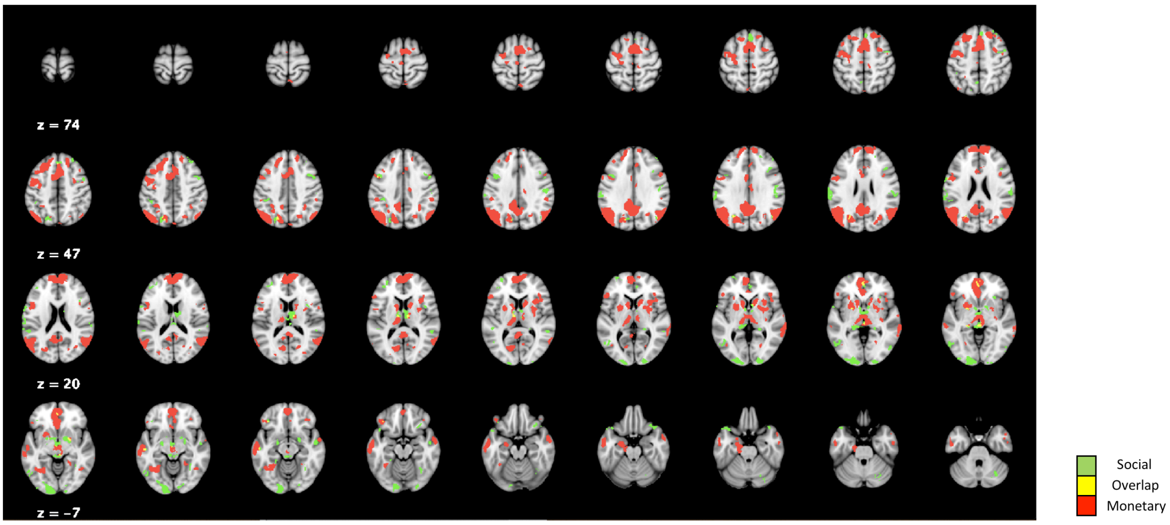
